# Ensemble viewing reshapes neural representations of same-race and other-race faces

**DOI:** 10.64898/2026.09.17.752481

**Authors:** Moaz Shoura, Han Ning Jiang, Zaynab Azeem, Otilia Iancu, Marco A. Sama, Jonathan S. Cant, Adrian Nestor

**Author notes:** Correspondence concerning this article should be addressed to Adrian Nestor, Department of Psychology at Scarborough, University of Toronto, 1265 Military Trail, Scarborough, Ontario, Canada, M1C 1A4.

## Abstract

The other-race effect, better recognition of same-race than other-race faces, is established for individual faces. Whether this benefit is reflected in neural representations of face ensembles, as when viewing a crowd, remains unclear. We recorded electroencephalography responses from 22 East Asian adults viewing synthetic East Asian (same-race) and White (other-race) faces in two formats: individually and in six-face ensembles. Multivariate decoding and generative image reconstruction assessed neural discriminability, representational geometry and recoverable visual content. Behavioral testing confirmed a same-race recognition advantage. The corresponding neural decoding advantage depended on viewing format and was not preserved for race-homogeneous ensembles, although their constituents retained it when viewed individually. Early ensemble decoding instead favored other-race faces. Race organized ensemble representations and reconstructed latent vectors matched ensemble summary targets above chance. Independent judgments revealed format-dependent race differences in reconstructed age and arousal relative to model-derived references. These findings constrain accounts of same-race expertise: its benefit for neural discriminability depends on whether faces are represented individually or collectively. Recoverable ensemble-summary information can also coexist with systematic appearance shifts, linking the organization of neural representations to their visual content.

## Introduction

Face recognition depends on representations that support individuation despite variation in appearance [1,2]. Experience with faces from one’s own racial group shapes these representations [3]. The resulting other-race effect (ORE), better recognition of same-race (SR) than other-race (OR) faces [4,5], varies with lifetime experience and opportunities for contact [6,7]. Understanding its neural basis requires characterizing both how distinctly faces are represented and what visual information their representations preserve. We therefore examine race-related differences in neural discriminability, geometry, temporal dynamics and recoverable content, without assuming that every measure must reflect the behavioral SR advantage.

Face-space accounts propose that identities occupy an experience-tuned multidimensional space whose dimensions capture facial variation relevant to distinguishing individuals [8]. Consistent with this principle, comparisons between face-recognition networks and human and monkey neurons link identity coding to representations of facial features [9]. These accounts predict greater separation among SR faces along identity-diagnostic dimensions, reflecting more extensive experience, whereas OR faces should occupy a denser region ([10,11,12]; but see [13]). Computational modeling, neural decoding and image reconstruction support aspects of this account [14,15,16]. Race-related differences in face processing also extend beyond identity recognition to expression and age judgments [17,18,19,20], while implicit racial attitudes can shape the visual information used to judge trustworthiness [21].

Most ORE evidence concerns faces viewed individually, whereas everyday encounters often involve groups, or ensembles. Observers rapidly estimate properties of face ensembles, including average identity and expression [22,23]. Ensemble representations can reduce noise and weight constituents unequally [24,25], but apparent averaging may also reflect subsampling, range information or interactions among summary statistics [26,27]. Here, *viewing format* distinguishes individual faces from six-face ensembles, with each ensemble treated as a stimulus in its own right. This comparison tests whether race-related differences in individual-face representations also characterize ensemble representations. Reconstruction further tests for information consistent with a model-defined ensemble mean, obtained by averaging constituent vectors in a relevant model’s latent space.

Accounts of SR expertise yield contrasting predictions for collective viewing. If ensemble discrimination depends on the fidelity of constituent face representations, greater SR precision should support an SR advantage in both viewing formats. Alternatively, if pooling downweights the individuating information that benefits from SR expertise, the SR advantage should diminish even while racial-composition information remains available. Behavioral studies report qualitatively different patterns: greater weighting of OR faces in racial-composition judgments [28], comparable SR and OR averaging [29,30], and selective discounting of OR information in emotion judgments [31]. Because these findings are inferred from different explicit judgments, their divergence could reflect differences in representations formed during viewing, their task-dependent use in decisions, or both. We therefore test whether viewing format changes the SR–OR decoding difference during an orthogonal task and examine how race shapes the temporal dynamics and recoverable content of ensemble representations.

Event-related potential (ERP) studies implicate early perceptual encoding in the ORE [32,33]. However, condition-averaged ERP component amplitudes can obscure trial-level heterogeneity [34] and do not directly characterize the discriminability or geometry of stimulus-specific response patterns. To address this, multivariate EEG and MEG analyses characterize face representations over time [2,35] and quantify their discriminability and representational structure [36,37]. For example, multivariate EEG detected changes in facial-attractiveness representations after social influence without corresponding reliable ERP effects [38]. Generative approaches further link visual appearance to neural representations [39,40,41], with image reconstruction making recoverable content available for direct comparison and independent judgment [42]. Successful matching of reconstructed latent vectors to their targets rather than foils can coexist with systematic appearance shifts. Evaluating age and affect can therefore reveal information that target-matching accuracy alone does not capture.

We combine these approaches to determine how viewing faces collectively changes race-related neural representations. East Asian adults viewed neutral, female-appearing East Asian (SR) and White (OR) faces individually and in ensembles during an orthogonal oddball task. A generative adversarial network (GAN), trained from scratch on equally sized race-by-sex image sets, generated the experimental face stimuli. Six-face ensembles comprised race-homogeneous sets (6:0), equal-split sets (3:3) and outlier sets (5:1). Four homogeneous, two equal-split and four outlier ensembles were tested with constituent positions varied across trials for each ensemble. Averaging the constituents’ GAN latent vectors defined a model-defined ensemble mean. The generator rendered this vector as a face image, providing an ensemble summary target distinct from a pixel average or an independently measured perceptual mean (Fig. 1).

**Figure 1.**
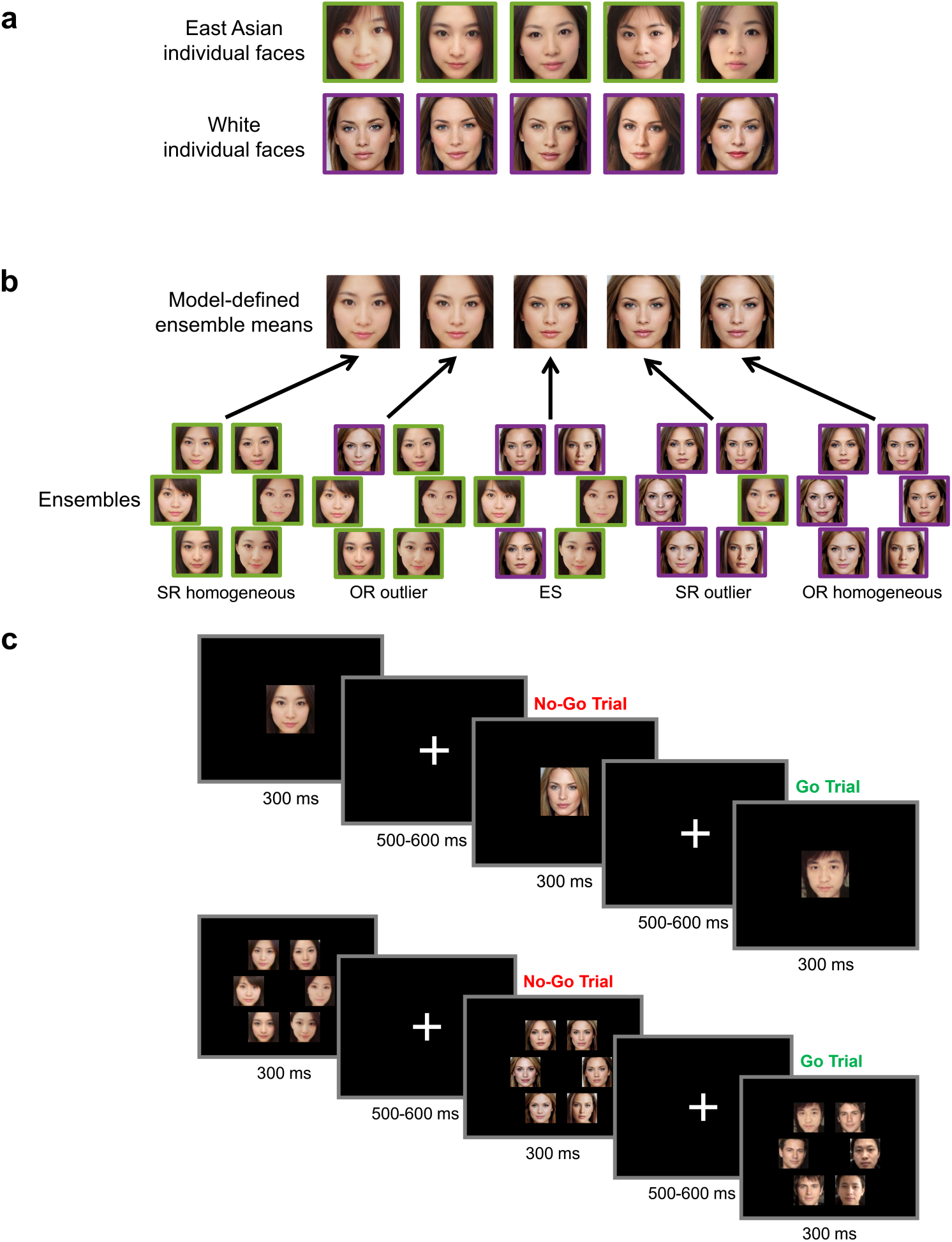
| Stimuli and procedure. **a** Examples of East Asian and White female-appearing faces. **b** Examples of six-face ensembles varying in racial composition: homogeneous ensembles contained six faces of one race, outlier ensembles contained five faces of one race and one of the other, and equal-split (ES) ensembles contained three faces of each race. Outlier labels identify the minority face. Images of the corresponding model-defined ensemble means, shown above, were generated by averaging the constituent latent vectors in StyleGAN2 *W* space and passing each mean vector through the generator. These summary images were also presented individually. **c** Participants responded to male-face oddballs, maintaining attention without an explicit identity, race or ensemble-summary judgment. All depicted faces are synthetic images generated using the study-specific StyleGAN2 model.

Individual-face responses provided a reference for comparing formats. For reconstruction, EEG dissimilarities positioned a stimulus in neural face space. Then, relationships between those coordinates and GAN features estimated a latent vector, which the generator rendered as a face image (Fig. 2).

**Figure 2.**
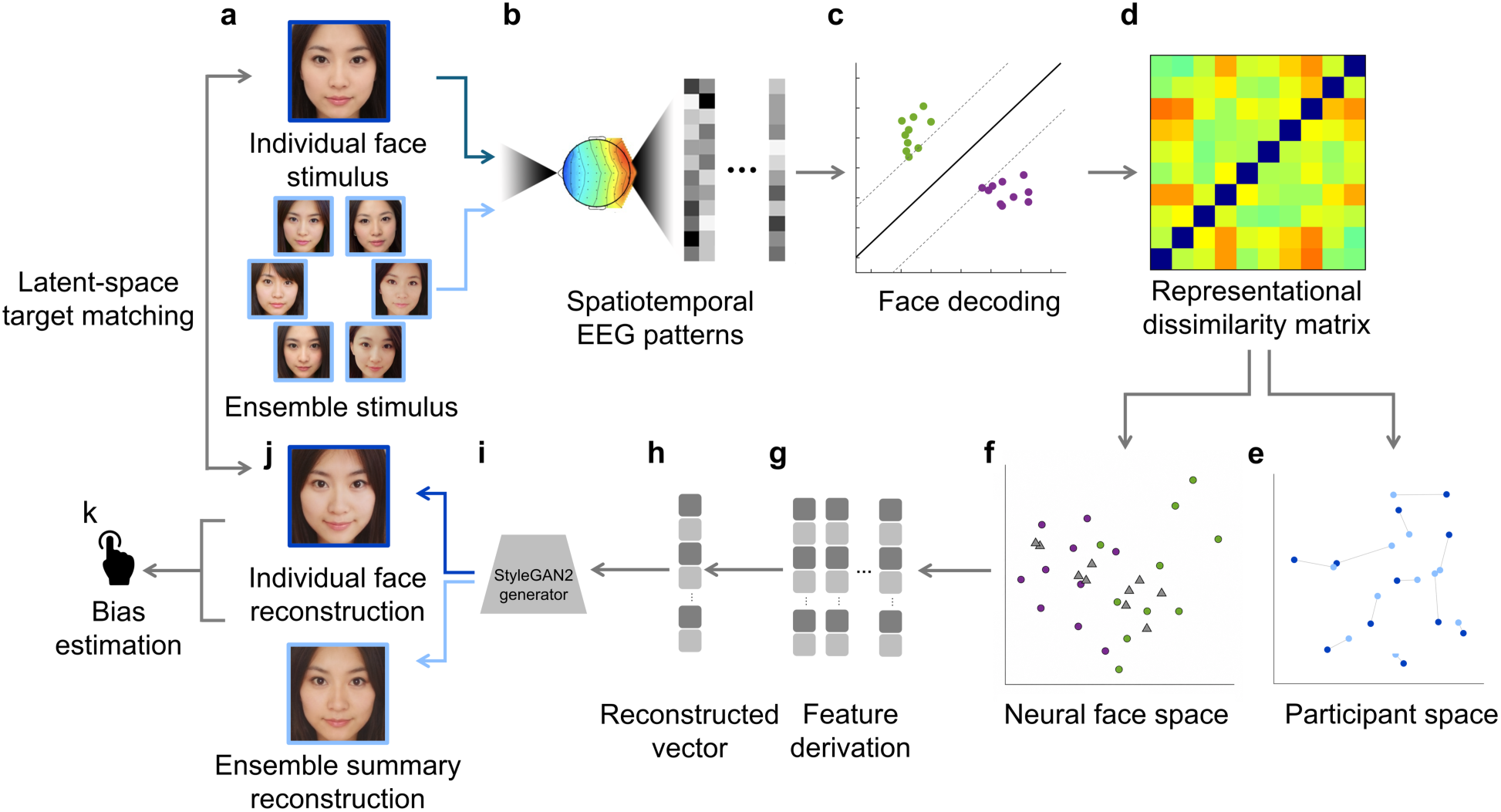
| From EEG patterns to representational geometry and reconstructed appearance. **a–d**, EEG responses to individual faces and six-face ensembles (**a**) provided spatiotemporal patterns (**b**) for pairwise linear classification (**c**); decoding accuracies were assembled into representational dissimilarity matrices (***d***). **e,f**, Principal component analysis represented participant-specific decoding profiles in participant space (**e**), whereas multidimensional scaling characterized relationships among stimuli in neural face space (**f**). **g–j**, Neural face-space coordinates were used to derive latent feature vectors (**g**), which were combined according to a held-out target’s coordinates to estimate its latent representation (**h**). The StyleGAN2 generator (**i**) transformed this vector into an individual-face or ensemble-summary reconstruction (**j**). Target-matching accuracy measured whether reconstructed latent vectors were closer to individual-face or ensemble summary targets than to held-out validation foils. ***k***, Independent observers compared EEG-based and theoretical-observer reconstructions to assess systematic differences in apparent age, valence and arousal. Face images are study-generated StyleGAN2 stimuli or EEG-based reconstructions (Methods and Supplementary Methods). The figure illustrates the analysis rather than numerical results.

Ensemble reconstructions were evaluated by matching reconstructed latent vectors to ensemble summary targets rather than validation foils. Independent observers then compared the age, valence and arousal of EEG-derived reconstructions with reference images reconstructed from the GAN’s own similarity structure. This design links discriminability to recoverable appearance without requiring age or affect to be specified as decoding targets.

Together, these analyses test whether the neural benefit associated with SR expertise depends on whether faces are represented individually or collectively. The individual-face SR decoding advantage was not preserved for homogeneous ensembles, even though their constituents retained that advantage when viewed individually. Reconstruction recovered information consistent with model-defined ensemble means, and independent judgments revealed format-dependent race differences in recovered age and arousal. This integrated characterization connects changes in discriminability to the summary information and appearance retained during collective viewing.

## Results

### Behavioral performance

Twenty-two East Asian participants viewed individual faces and six-face ensembles (Fig. 1a,b) during EEG recording. Sensitivity in the orthogonal male-oddball task was near ceiling (*d*′ = 4.997; Fig. 1c). Participants performed better on the Chinese (SR; [43]) than White (OR; [44]) Cambridge Face Memory Test (CFMT; *t*(21) = 7.93, *p* < .001, *d* = 1.69), establishing a robust behavioral ORE in this sample.

### Viewing format changes the same-race decoding advantage

Linear pairwise decoding used leave-one-block-out cross-validation to classify every stimulus pair for individual faces and ensembles (Fig. 2). Spatiotemporal decoding concatenated EEG amplitudes across 12 occipitotemporal (OT) electrodes and all time points from 50 to 650 ms into one feature vector per observation, retaining information across early and later face processing [45,46]. We refer to this as *broad-window decoding*. Accuracies were summarized for SR, OR and between-race individual-face pairs and for homogeneous, equal-split (ES) and outlier ensemble comparisons. An all-electrode analysis provided a complementary check (Supplementary Fig. 1).

For individual faces, decoding accuracy was significantly above chance for SR, OR, and between-race comparisons (Wilcoxon signed-rank tests; all *ps* < .001; Fig. 3a). Accuracy differed across the three classification types (*F*(2, 42) = 58.52, *p* < .001, *ηₚ²* = .736), with between-race face pairs decoded more accurately than both SR (*t*(21) = 4.15, *p* = .001, *d* = 0.469) and OR pairs (*t*(21) = 10.79, *p* < .001, *d* = 1.540). As predicted, SR faces were decoded more accurately than OR faces (*t*(21) = 6.10, *p* < .001, *d* = 1.070), consistent with prior evidence of race-related differences in neural discriminability [15]. The all-electrode analysis yielded the same pattern at generally lower accuracy (Supplementary Fig. 1).

**Figure 3.**
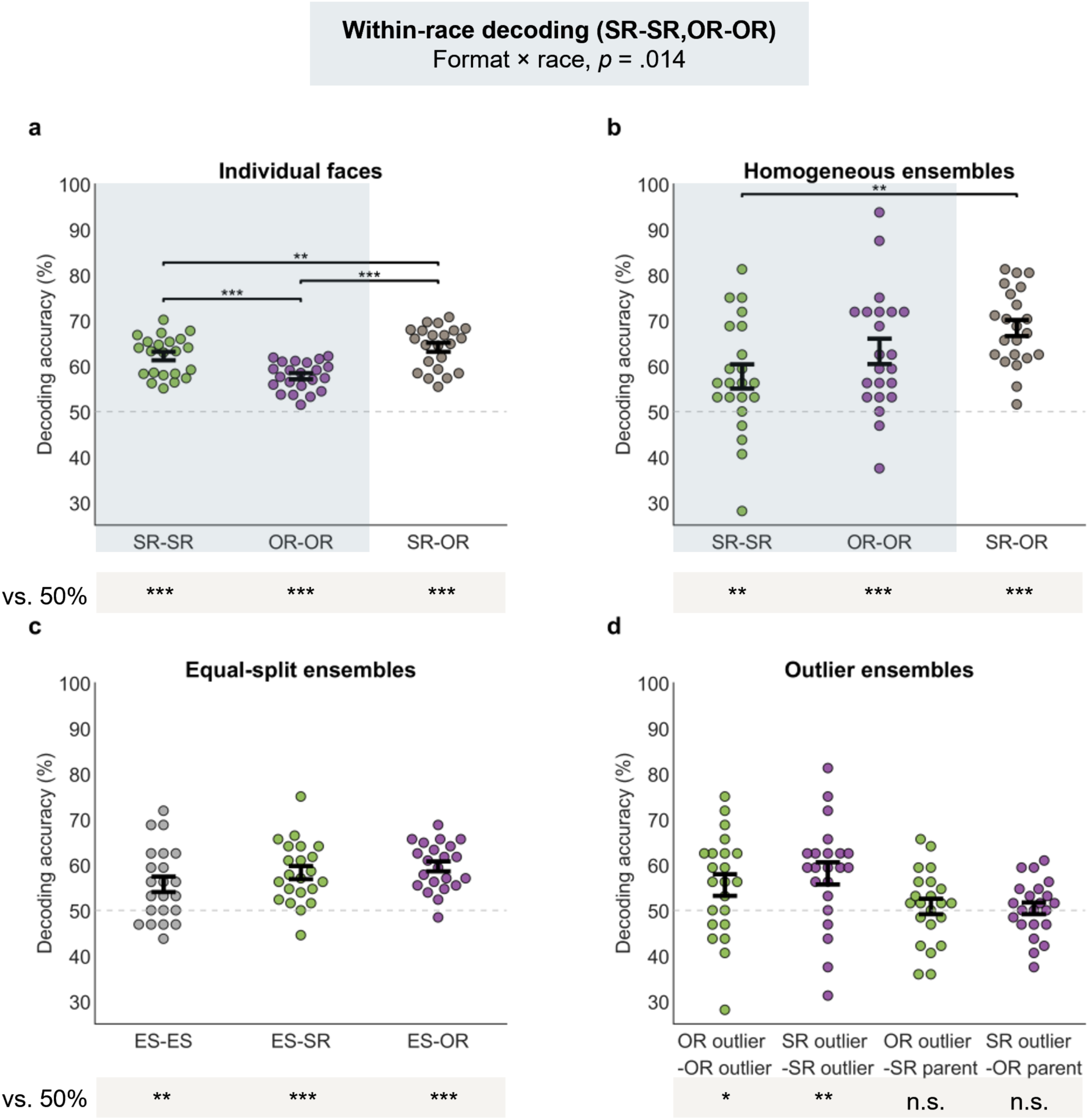
| Ensemble viewing changes the same-race advantage in neural discriminability. Spatiotemporal decoding concatenated EEG activity across 12 occipitotemporal electrodes and 50–650 ms. **a**, Individual-face decoding was highest between races and higher within same-race (SR) than other-race (OR) faces. **b**, Homogeneous ensembles were more discriminable between races than within SR ensembles, without a reliable SR–OR difference. Pale-blue backgrounds in (**a)** and (**b)** identify the within-race categories entered into the viewing-format × race analysis; between-race decoding was excluded. **c**, Equal-split (ES) ensembles were discriminable from one another and from homogeneous ensembles, without reliable differences among comparison types. ***d***, Outlier ensembles were discriminable from one another but not reliably from their homogeneous parents, with which they shared five of six faces. Outlier labels identify the minority face. Dots represent participants (*n* = 22). Error bars indicate ± SEM. Dashed lines mark 50% chance. Brackets indicate two-sided post hoc contrasts with Bonferroni correction across three comparisons per panel in **(a)** and **(b)**. Grey strips show two-sided Wilcoxon signed-rank tests against chance, with Bonferroni correction across the three tests. \**p* < .05, \*\**p* < .01, \*\*\**p* < .001; *n*.s., not significant.

Homogeneous ensembles were decoded above chance within and between races (Fig. 3b). Decoding differed across the three comparison types (*F*(2, 42) = 5.64, *p* = .007, ηₚ² = .212). Between-race ensemble accuracy was higher than SR ensemble accuracy (*t*(21) = 3.54, *p* = .006, *d* = 0.924), whereas SR and OR homogeneous ensemble decoding did not differ reliably (*p* = .455). The all-electrode analysis showed a similar pattern (Supplementary Fig. 1).

To test directly whether the SR–OR decoding difference varied with viewing format, accuracies were entered into a 2 × 2 repeated-measures ANOVA with viewing format (individual face, homogeneous ensemble) and race (SR, OR) as within-participant factors. Neither format (*F*(1, 21) = 0.063, *p* = .805) nor race (*F*(1, 21) = 0.084, *p* = .774) had a significant main effect. Critically, the interaction was significant (*F*(1, 21) = 7.115, *p* = .014, ηₚ² = .253; Fig. 3a,b). Thus, viewing format changed the SR–OR decoding difference: the individual-face SR advantage was not preserved for homogeneous ensembles.

To assess whether this interaction reflected an atypical constituent sample, we restricted the individual-face decoding analysis to the faces used in homogeneous ensembles. SR constituents were more discriminable than OR constituents both for pairs drawn from different ensembles of the same race (*t*(21) = 3.036, *p* = .003, *d* = 0.65; Supplementary Fig. 2) and for pairs within the same ensemble (*t*(21) = 1.873, *p* = .038, *d* = 0.40; both tests one-tailed and unadjusted). These controls show that the selected constituents retained an SR decoding advantage when viewed individually, arguing against the ensemble result arising simply because its constituents lacked this advantage.

Additional analyses examined ES and outlier ensembles. ES ensembles were decoded above chance from one another and from homogeneous ensembles, without reliable differences across ES–ES, ES–SR, and ES–OR comparisons (one-way repeated-measures ANOVA, *p* = .134; Fig. 3c). Outlier ensembles were also discriminable from one another, but outlier–parent decoding did not exceed chance (both *ps* ≥ .623), with these displays sharing five of six constituents. Decoding accuracy did not differ reliably between the two racial compositions, either for pairs of outlier ensembles with the same racial composition (*p* = .479) or for outlier–parent pairs (*p* = .600; Wilcoxon signed-rank tests; Fig. 3d).

Race therefore structured neural discriminability in both formats, while the interaction and constituent controls show that the individual-face SR advantage did not transfer unchanged to collective viewing. We next examined how these decoding relationships were organized in neural face space.

### Race organizes individual-face and ensemble geometry

Pairwise decoding accuracies were submitted to multidimensional scaling (MDS), separately for each participant and format. Retaining at least 90% of the variance required 24-25 dimensions for individual faces and 5-6 dimensions for ensembles. The different numbers of stimuli (64 individual faces versus 10 ensembles) limit direct comparison of these dimensionalities.

Individual-face neural spaces showed clear separation by race, with OR faces occupying a more compact region than SR faces (Fig. 4a). Across participants, distances between SR faces were larger than distances between OR faces (Wilcoxon signed-rank tests, median difference Mdn_diff_ = 3.72%; *W* = 243, *p* < .001) whereas between-race distances exceeded both OR (Mdn_diff_ = 4.78%, *W* = 253, *p* < .001) and SR distances (Mdn_diff_ = 0.98%, *W* = 228, *p* < .001). By embedding decoding-derived dissimilarities in a Euclidean space, this analysis quantifies SR–OR differences in representational dispersion, linking neural discriminability to the organization predicted by face-space accounts.

**Figure 4.**
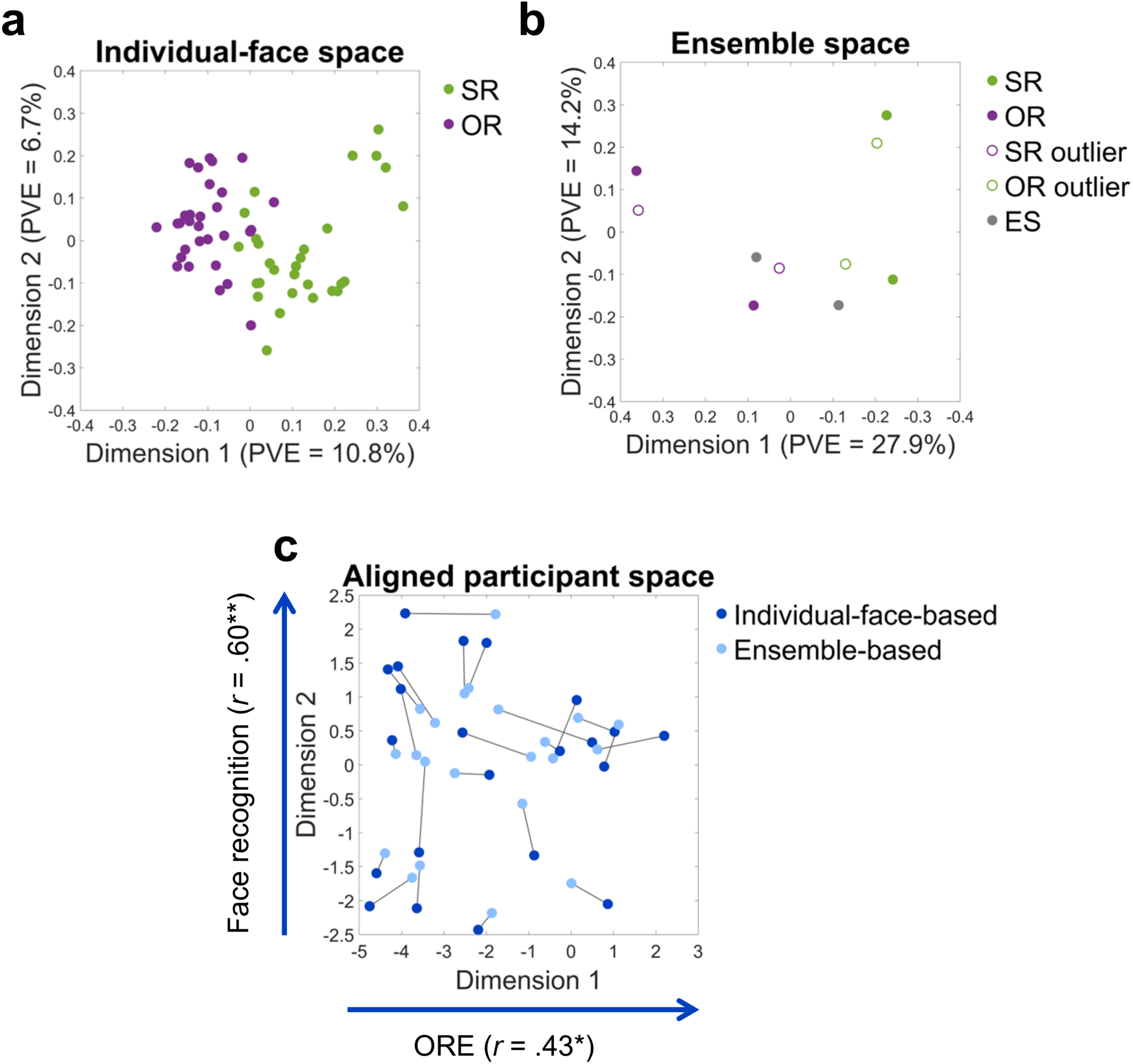
| Race organizes neural face spaces, while individual differences link decoding to recognition. **a,b**, Two-dimensional multidimensional scaling visualizations derived from participant-averaged decoding dissimilarities for 64 individual faces (**a**) and 10 ensembles (**b**). Each point represents one stimulus; axis labels indicate the percentage of variance explained (PVE). Other-race (OR) individual faces occupied a more compact region than same-race (SR) faces. Ensembles were organized by racial composition, with equal-split (ES) displays between homogeneous displays and outlier displays near their homogeneous parents. Open markers identify outlier ensembles; their labels specify the minority face’s race and their colors indicate the majority race. **c**, Participant configurations obtained separately by principal component analysis (PCA) and aligned by Procrustes analysis. Lines connect the same participant across formats; axes follow the individual-face reference configuration. Arrows indicate associations between individual-face PC1 scores and the behavioral other-race effect (ORE; SR minus OR Cambridge Face Memory Test performance; Pearson’s *r* = .430, *p* = .046), and between PC2 scores and SR CFMT performance (*r* = .601, *p* = .003). Correlation tests were two-sided and unadjusted; \**p* < .05, \*\**p* < .01. No corresponding ensemble associations were significant. Cross-format alignment exceeded chance correspondence (10,000 label permutations, *p* < .001). *n* = 22 participants.

Ensemble-derived neural spaces were also organized by racial composition (Fig. 4b). Homogeneous East Asian and White ensembles occupied distinct regions, ES ensembles lay between them, and outlier ensembles were close to their parent homogeneous ensembles. This descriptive geometry indicates that race continues to organize ensemble representations even though homogeneous SR and OR ensembles did not differ reliably in broad-window decoding accuracy.

As a supporting analysis, PCA mapped participants’ decoding profiles into a separate participant space for each viewing format (Fig. 4c). Following prior findings [15], scores on the first two components were related to SR CFMT performance and behavioral ORE magnitude (SR minus OR CFMT scores). For individual faces, PC1 correlated with ORE magnitude (*r*(20) = .430, *p* = .046) and PC2 with SR CFMT performance (*r*(20) = .601, *p* = .003; both uncorrected). No corresponding ensemble associations were reliable. Participant spaces nevertheless aligned across formats more closely than expected by chance (10,000 label permutations, *p* < .001), indicating shared structure in participant-specific decoding profiles. Additional analyses are reported in Supplementary Results.

Together, these analyses characterize individual and ensemble representations within a common face-space framework. The individual-face neural space additionally provides the reference for reconstructing both individual faces and ensemble summaries, linking neural discriminability to recoverable appearance.

### Ensembles show an early other-race decoding advantage

We next examined whether the SR advantage for individual faces and the altered pattern observed for ensembles reflected brief processing stages or information distributed across a broader temporal interval. Pairwise decoding was repeated in successive ∼20 ms sliding windows, with cluster-based correction across time.

For individual faces, decoding exceeded chance for both SR and OR faces from approximately 130 ms, peaked between 220 and 250 ms, and declined after approximately 650 ms (Fig. 5a). SR decoding was numerically higher over much of this period, although the fully time-resolved SR-OR difference did not survive cluster correction. The broad-window SR advantage is consistent with information distributed across time that becomes detectable when combined [15].

**Figure 5.**
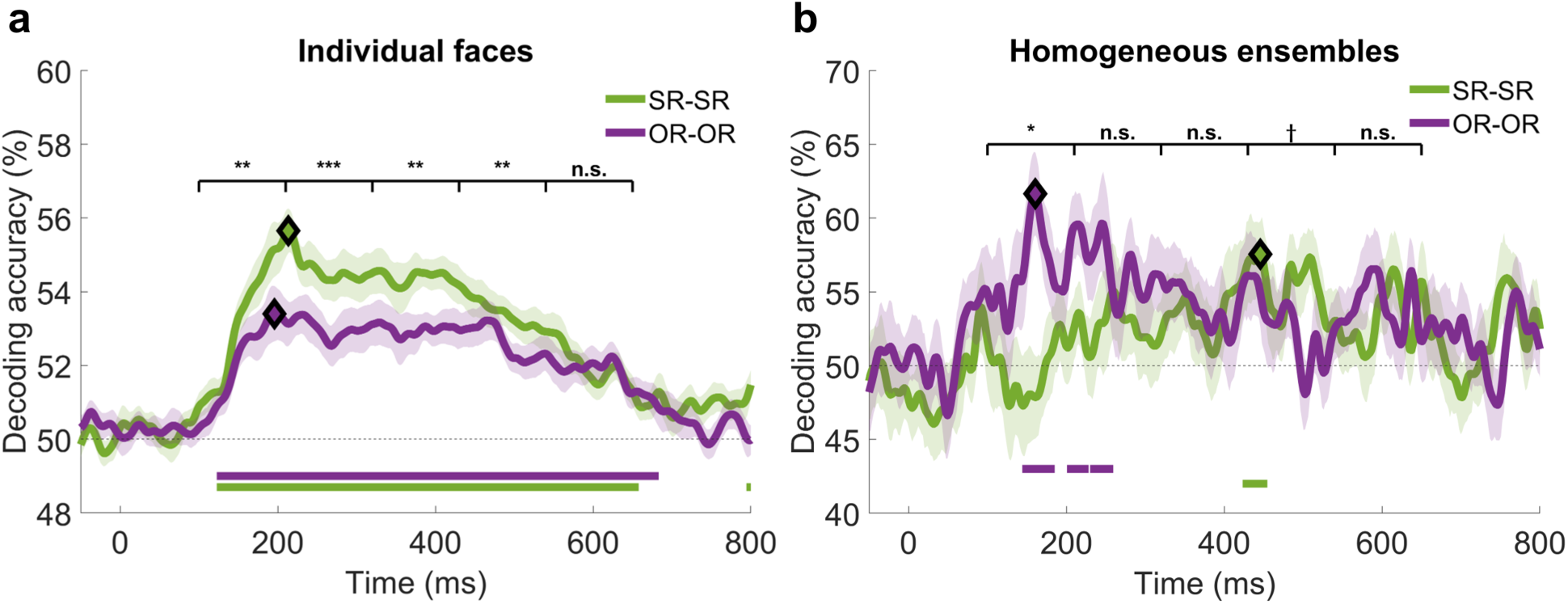
| Individual faces and homogeneous ensembles show different temporal profiles of within-race decoding. **a,b**, Decoding accuracy for pairs of same-race (SR) or other-race (OR) individual faces (**a**) and homogeneous ensembles (**b**) in successive approximately 20-ms windows. Lines show participant means, shading indicates ± SEM (standard error of the mean), and diamonds identify peaks in the group-average curves. Dashed lines indicate 50% chance accuracy; colored horizontal segments mark above-chance intervals after cluster-based correction across time (*p* < .05). Brackets summarize SR–OR comparisons using activity combined within five consecutive 110-ms windows from 100 to 650 ms. Individual-face decoding favored SR stimuli in the first four windows, whereas ensemble decoding favored OR stimuli in the earliest window. Fully time-resolved analysis relied on two-sided sign-flip permutation *t* tests (10,000 permutations). Window-wise comparisons used two-sided Wilcoxon signed-rank tests with Bonferroni correction across five windows: \**p* < .05, \*\**p* < .01, \*\*\**p* < .001; n.s., not significant; † *p* < .05, uncorrected. Fully time-resolved SR–OR contrasts yielded no significant clusters. *n* = 22 participants.

At an intermediate level of temporal resolution, we divided the 100–650 ms interval into five consecutive windows and concatenated activity across sensors and time points within each window for decoding. Decoding showed a significant SR advantage in the first four windows (100–210, 210–320, 320–430 and 430–540 ms; Bonferroni-corrected *p*s ≤ .02), but not in the final window (540–650 ms; *p* = .71). These results identify multiple intervals of greater SR discriminability, consistent with the main broad-window analysis.

Homogeneous ensembles showed a different temporal profile (Fig. 5b). OR decoding significantly exceeded chance from approximately 150 to 250 ms, whereas SR decoding exceeded chance from approximately 400 to 450 ms. The fully time-resolved SR–OR comparison was not significant after cluster correction; however, the intermediate-window analysis found a significant OR advantage during the earliest 100-210 ms interval (*p* = .02, Bonferroni-corrected across five windows). A later numerical SR advantage between 430-540 ms did not survive correction (*p* = .049, uncorrected).

For ES and outlier ensembles, only SR-outlier decoding exceeded chance over a significant interval (∼150–250 ms; Supplementary Fig. 3). Lower discriminability and information distributed across longer intervals may explain the weaker time-resolved results associated with these ensembles.

Together, these results reveal different temporal profiles of race-related neural discriminability: multiple decoding windows favored SR individual faces, whereas the earliest intermediate window favored OR homogeneous ensembles. This early OR advantage identifies a temporally specific feature of ensemble processing that overall decoding accuracy alone does not capture, highlighting how collective viewing reshapes the dynamics of neural face discrimination.

Sensor-level searchlights provided a complementary check: the SR advantage for individual faces was widespread and extended beyond the OT electrode set used for primary decoding. Ensemble decoding exceeded chance in several sensor neighborhoods, but no SR–OR ensemble contrast survived correction (Supplementary Fig. 4; Supplementary Results and Methods).

### Reconstruction recovers ensemble summary information across racial compositions

Decoding indicates whether patterns are separable but not what visual information they preserve. We therefore reconstructed individual-face and ensemble representations as vectors in StyleGAN2 *W* space from each participant’s decoding results (Fig. 2).

Briefly, dimensions of individual-face neural space were related to GAN latent features, which were combined according to the target’s projected coordinates and rendered as an image. The target’s latent vector was excluded from feature derivation. Ensemble summary representations were projected into the individual-face space using cross-format decoding of ensembles from individual faces (see Methods).

Target-matching accuracy measured whether a reconstructed latent vector was closer in *W* space to its target than to a validation foil (50% chance; same-race foils for individual faces). A theoretical observer (TO), replacing neural dissimilarities with *W*-space distances, provided a model-derived reference. TO reconstructions exceeded chance for both races (all *Ws* ≥ 428, all *p*s < .001; Wilcoxon signed-rank tests across reconstructions), without a significant SR–OR difference (*W* = 154, *p* = .186; Fig. 6a). This comparison documents the model reference for a stimulus set selected to avoid a significant TO race difference (Supplementary Methods).

**Figure 6.**
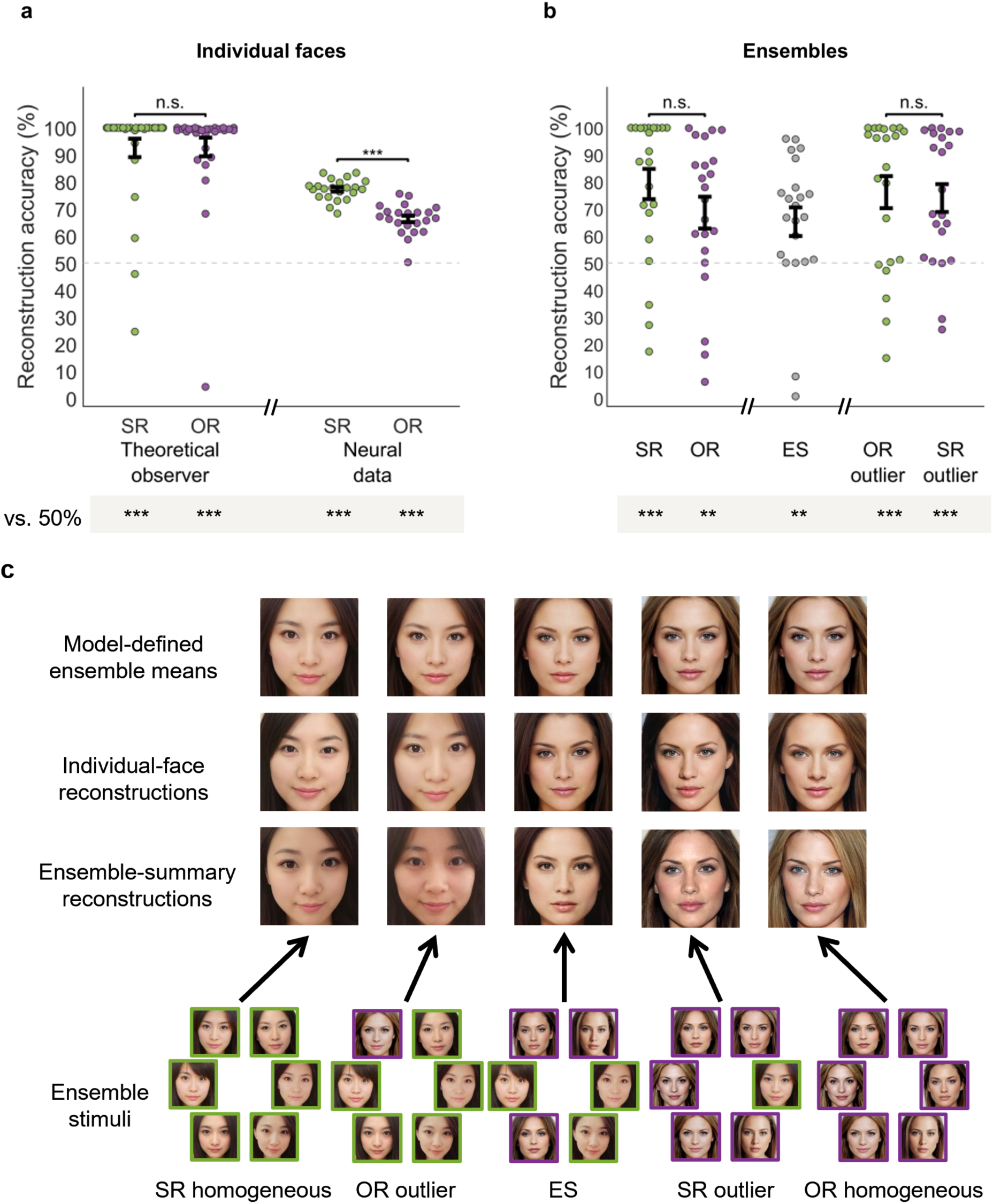
| Neural reconstruction recovers individual-face and ensemble-summary information. **a**, Individual-face reconstructions matched their targets above chance using either a theoretical observer (TO), based on generative adversarial network (GAN) latent-space distances, or neural data. Neural reconstruction showed a same-race (SR) advantage over other-race (OR) faces; the TO comparison was not significant. **b**, Neural reconstructions matched ensemble summary targets above chance for homogeneous, equal-split (ES) and outlier ensembles. Homogeneous SR and OR ensembles did not differ reliably, nor did the two outlier types. Across individual-face and homogeneous-ensemble viewing, a repeated-measures ANOVA identified an overall SR advantage (*p* = .025), without a format × race interaction (*p* = .969). Accuracy is the proportion of comparisons in which a reconstructed latent vector was closer to its target than to a validation foil; ensemble foils matched the target’s racial composition. Dots represent reconstructed target faces for the TO and participants for neural data (*n* = 22); black marks and error bars show means ± SEM (standard error of the mean). Dashed lines mark 50% chance; grey strips show comparisons against chance and brackets show between-condition comparisons. Neural comparisons used two-sided Wilcoxon signed-rank tests. **c**, Representative examples show model-defined ensemble means, their EEG-based reconstructions when viewed individually, and reconstructions elicited by the corresponding ensembles shown below. Outlier labels identify the minority face. All face images are study-generated StyleGAN2 stimuli or EEG-based reconstructions (Methods and Supplementary Methods). \*\**p* < .01, \*\*\**p* < .001; *n*.s., not significant.

EEG-derived reconstructions were also reliably above chance for both SR and OR faces across participants (all *Ws* ≥ 253, all *p*s < .001; Fig. 6a) – for examples, see Fig. 6c. Critically, reconstruction accuracy was higher for SR than OR faces (*W* = 252, *p* < .001).

We next reconstructed ensemble representations from each ensemble’s decoding profile relative to individual-face stimuli – Fig. 6c compares model-defined ensemble means, reconstructions elicited by those mean images viewed as individual faces, and reconstructions elicited by the corresponding ensembles. Reconstructions matched ensemble summary targets above chance for every racial composition, using composition-matched validation foils (all *Ws* ≥ 210, all *p*s < .007; Fig. 6b). Neither the homogeneous SR-OR contrast (*p* = .31) nor the SR-OR outlier contrast (*p* = .61) was significant.

A 2 × 2 repeated-measures ANOVA across individual face and homogeneous ensemble reconstructions showed a main effect of race (*F*(1, 21) = 5.785, *p* = .025, *η*^2^ = .216), but not of format (*F*(1, 21) = 0.265, *p* = .612, *η*^2^ = .012) or interaction (*F*(1, 21) = 0.002, *p* = .969, *η*^2^ < .001). Thus, reconstruction supported an overall SR advantage without evidence that its magnitude changed with viewing format.

### Recovered appearance reveals systematic differences beyond target matching

Successful target matching in GAN latent space does not preclude systematic shifts in reconstructed appearance. Building on prior age and expressiveness findings [15,16], we therefore examined the age, valence, and arousal conveyed by the reconstructions.

Forty-four observers, two per EEG participant’s reconstruction set, judged each EEG reconstruction against its TO counterpart. Here, bias denotes a directional judgment difference from this model-derived reference, with 50% indicating no preference.

SR reconstructions appeared older than their TO counterparts and OR reconstructions younger, both for individual faces and homogeneous ensembles (all FDR-adjusted *ps* ≤ .0363; Fig. 7a). An aligned rank transform (ART) ANOVA revealed a race effect (*F*(1, 21) = 25.25, *p* < .001) and an interaction with viewing format (*F*(1, 21) = 10.43, *p* = .004), with a descriptively larger SR–OR separation for ensembles. ART-C comparisons confirmed SR–OR differences for individual faces (*p* = .010) and homogeneous ensembles (*p* = .002).

**Figure 7.**
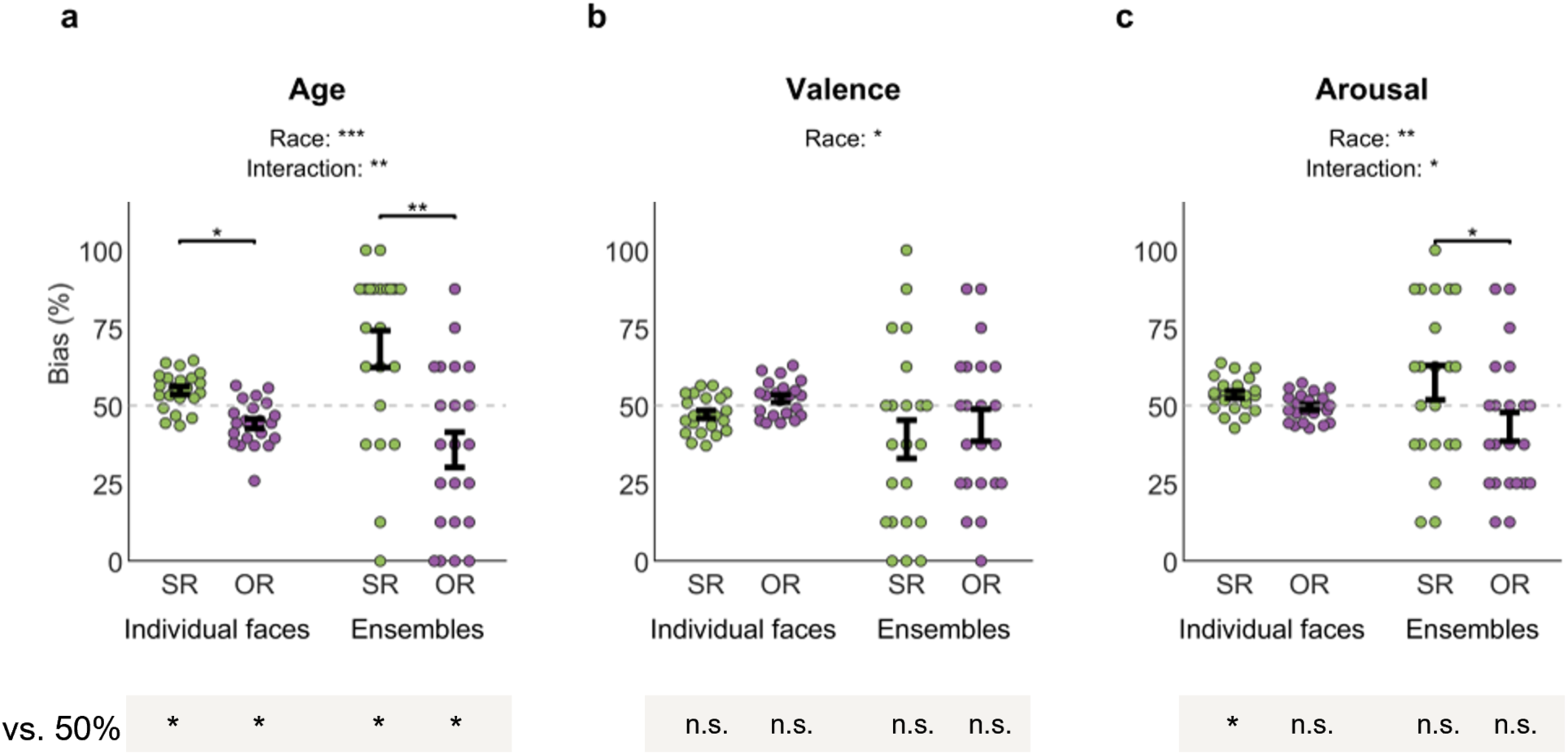
| Race and viewing format shape reconstructed appearance. Observers compared EEG-derived reconstructions with theoretical-observer (TO) references. Bias (%) indicates how often EEG reconstructions appeared older (a), more positive (b), or more emotionally intense (c); the dashed line marks 50%, indicating no directional preference. SR, same-race (East Asian); OR, other-race (White). Race effects occurred for all attributes, with race-by-format interactions for age and arousal. Relative to TO references, SR reconstructions appeared older and OR reconstructions younger in both formats; only SR individual-face reconstructions showed elevated arousal, and no valence condition differed significantly. Dots show means for 22 reconstruction sets, each averaging two observers; black marks show the grand mean ± SEM across sets. Mixed-effects aligned rank transform (ART) ANOVAs accounted for observer and reconstruction-set grouping. Brackets show significant ART-C contrasts, with FDR correction across six comparisons per attribute. Two-sided clustered Wilcoxon tests against 50% were FDR-corrected across 12 conditions. Omnibus *p* values are unadjusted; other *p* values are FDR-adjusted. \**p* < .05, \*\**p* < .01, \*\*\**p* < .001; *n*.s., not significant.

Reconstructed affect showed a different pattern. Model-relative valence judgments were lower for SR than OR reconstructions (*F*(1, 21) = 5.26, *p* = .032), although no condition differed significantly from its TO reference (all *ps* > .13; Fig. 7b). The race difference did not vary reliably with viewing format (*p* = .649).

Arousal judgments were higher for SR than OR reconstructions (*F*(1, 21) = 10.92, *p* = .003), and this difference varied with viewing format (*F*(1, 21) = 4.81, *p* = .040). ART-C comparisons identified an SR–OR difference for homogeneous ensembles (*p* = .031), but not individual faces (*p* = .127). Relative to the TO reference, only SR individual-face reconstructions appeared more emotionally intense (*p* = .014; all other *ps* ≥ .207; Fig. 7c). No main effect of viewing format was detected for any attribute (all *ps* > .13).

Together, these findings reveal opposing age shifts for SR and OR reconstructions, with race differences in age and arousal varying across viewing formats. These appearance judgments complement latent-space target matching by characterizing systematic shifts in the visual content recovered through reconstruction.

## Discussion

Our findings identify viewing format as a constraint on the neural advantage associated with SR expertise. The individual-face SR decoding advantage was not preserved for homogeneous ensembles, although race continued to organize the neural geometry of both formats. Reconstruction recovered ensemble-summary information across racial compositions, while independent judgments revealed format-dependent race differences in reconstructed age and arousal. These findings connect neural discriminability with recoverable appearance, showing why neither measure alone fully characterizes race-related face representations.

### Viewing format constrains the same-race advantage

A straightforward transfer of SR expertise to collective viewing would predict an SR ensemble advantage inherited from more precise constituent representations. Instead, the decoding interaction is consistent with pooling reducing the influence of the individuating information that benefits from SR expertise [47,48], while retaining summary properties such as average appearance and racial composition [24,25]. Crucially, the selected ensemble constituents retained an SR decoding advantage when viewed individually, arguing against stimulus selection alone as an explanation for the format effect.

For individual faces, the OR decoding disadvantage and denser OR geometry are consistent with experience-tuned face-space accounts [3,8,10]. By embedding pairwise discriminability in a common Euclidean space, we characterized the relative positions and dispersion of face representations, connecting decoding patterns to a geometric account of individuation. Agreement across OT, all-electrode and searchlight analyses supports the individual-face SR advantage. Further, results with female-appearing faces complement earlier decoding and reconstruction findings obtained with male-appearing faces [15,16], broadening the evidence across stimulus sets.

These findings also support a perceptual contribution to the ORE. Viewing format modulated SR–OR differences in neural discriminability without explicit identity or race judgments, implicating differences in the representations available during perception. This interpretation complements evidence from simultaneous face matching [49] and early electrophysiological responses [32,33] regarding a perceptual component alongside the established recognition-memory disadvantage for OR faces [5,33].

Supporting analyses related participant-space configurations derived from individual-face decoding to SR recognition performance and ORE magnitude across different stimuli and tasks, complementing prior findings [15]. These behavioral measures capture distinct aspects of performance: reduced recognition of both SR and OR faces can coexist with a preserved ORE [50]. Participant configurations also corresponded across viewing formats, although the ensemble associations with behavior were not reliable. Larger samples could establish how recognition ability and race-related advantages contribute to this shared neural decoding structure.

Temporal analyses further distinguish the individual-face and ensemble profiles. Homogeneous SR ensembles showed late above-chance decoding around 400–450 ms, consistent with the 408 ms peak reported for White observers viewing White-face ensembles [46]. This correspondence suggests broader generality of late SR ensemble discriminability across observer groups and synthetic versus photographic stimuli. Homogeneous OR ensembles instead showed early above-chance decoding and an OR advantage in the 100–210 ms window, contrasting with the individual-face SR advantage across windows spanning 100–540 ms. These SR–OR differences emerged when information was combined within intermediate-size windows while neither fully time-resolved contrast survived correction. Viewing format thus shaped the timing and direction of detectable race differences, highlighting the value of examining information at multiple temporal scales.

The early ensemble effect invites comparison with the other-race categorization advantage (ORCA; [51,52]), whereby OR faces can be categorized by race more efficiently than SR faces. One possibility is that OR ensembles were separated early using coarse but readily accessible features, overlapping with category-relevant ones. This interpretation draws on individual-face accounts emphasizing categorization for OR faces versus SR individuation [47,53] while our decoding distinguished ensembles within each race rather than race categories. Comparing neural dissimilarities with multiple models of image properties and facial structure across observer groups viewing identical stimuli could help determine whether the early advantage reflects stimulus differences, experience-dependent feature weighting, or their combination.

### Ensemble summaries remain recoverable across racial compositions

Ensemble geometry provides a bridge from discriminability to recoverable summary information. Homogeneous East Asian and White displays occupied separate regions in neural face space, with ES ensembles between them and outlier ensembles near their parents. This organization is consistent with summaries that retain racial-composition information despite the altered within-race decoding difference [22,27]. Outlier–parent proximity is compatible with reduced outlier weighting [24,31], for displays sharing five of six constituents. However, the present data therefore does not conclusively distinguish complete outlier exclusion or partial downweighting from equal-weight pooling.

Reconstruction further links neural face-space geometry to recoverable visual content. EEG-derived individual-face reconstructions supported above-chance target matching, with higher accuracy for SR than OR faces, building on previous neural and behavioral reconstruction work [15,16]. All ensemble types also supported above-chance matching to ensemble summary targets across racial compositions. Recovering this summary information through the individual-face neural embedding places both formats within a common representational framework. Reconstruction accuracy showed an overall SR advantage without a reliable format interaction, so the decoding interaction should not be generalized to all measures of recoverable information.

### Race and viewing format shape reconstructed appearance

Beyond target matching, data-driven reconstruction makes representational content available for visual and behavioral assessment [42]. Reconstructions have, for example, captured attention-related changes in apparent image contrast [54]. Here, independent judgments revealed model-relative shifts in reconstructed age and affect, although these attributes were not specified as decoding targets. Generative models thus provide experimental tools for investigating biases in human representations against an explicit model-derived reference, complementing generative analyses of culturally variable expression judgments [55] and broader proposals for their use in cognitive neuroscience [56]. Successful target matching can coexist with systematic appearance shifts; evaluating recovered attributes therefore adds information that reconstruction accuracy alone cannot provide.

For age, prior work showed race-dependent estimation accuracy [17,19], and image-based reconstruction, which did not rely on a GAN, revealed younger-looking OR than SR representations [15]. The present findings converge with this relative pattern and additionally identify opposing shifts from corresponding TO references in both viewing formats: SR reconstructions appeared older and OR reconstructions younger. Regarding the larger SR–OR separation in ensemble judgments, one possibility is that it reflects pooling that reduces identity-specific variability while preserving shared age-related cues, making the opposing judgments more consistent. These shifts concern model-relative appearance rather than chronological age and do not establish a universal OR-younger bias: for instance, explicit judgments have also overestimated the age of White faces [19].

Race also affects expression categorization and perceived emotional intensity [18,20,57]. Our findings connect this literature to the affective appearance recovered from neural responses to neutral faces: SR reconstructions received lower model-relative valence but higher arousal judgments than OR reconstructions. This combination could reflect race-dependent weighting of subtle affective cues. The greater SR–OR arousal separation for ensembles further suggests that pooling may change how cues to emotional intensity contribute to reconstructed appearance, complementing evidence that racial composition affects perceived crowd emotionality [58]. Such appearance differences remained detectable even without a reliable SR ensemble decoding advantage, reinforcing the value of assessing recovered content alongside discriminability.

### Generative reconstruction enables broader tests of perception

Generative reconstruction offers a flexible way to examine dimensions of appearance that were not specified during decoding. The differing patterns for age, valence and arousal motivate further tests of how facial dimensions are preserved or transformed during collective viewing [26,59]. The same reconstructed images could support judgments of trustworthiness or competence without fitting a separate neural decoder for each attribute, within the range of appearances supported by the generator. Prior evidence of race-related differences in the visual cues supporting trustworthiness judgments [21] makes this a particularly relevant application.

This approach could also test how predictive context shapes recovered appearance. Learned scene–face associations can evoke anticipatory face representations [60], motivating comparisons of identical individual faces and ensembles following different predictive cues. Holding the viewed stimuli constant would test whether expectations shift reconstructed age or affect, whether these shifts depend on viewing format, and whether target matching remains intact.

### Limitations and future directions

Several limitations define the scope of these findings. All EEG participants were East Asian; reciprocal observer samples would help distinguish effects of observer–face racial correspondence from stimulus-category differences, particularly given cultural differences in ensemble weighting [61]. Ensemble comparisons also used limited numbers of stimuli from a constrained synthetic stimulus set, motivating replication across more identities and ensembles. Finally, StyleGAN2 provides a controlled generative framework for stimulus design and reconstruction [16], but its geometry and the theoretical observer remain model-dependent reference structures despite balanced training.

## Conclusions

These findings indicate that the advantages associated with SR expertise depend on whether faces are viewed individually or collectively. Ensemble viewing altered the SR–OR decoding difference while preserving recoverable information and revealing systematic differences in reconstructed appearance. The neural differences observed without explicit identity or race judgments support a perceptual representational contribution to the ORE. Accounts of race-related face processing should therefore explain both how well neural representations can be discriminated and what visual information they retain.

## Methods

### Participants

A total of 22 East Asian adults (age range: 18-27 years; 12 women) from the University of Toronto community completed the EEG experiment. All participants were born and raised in East Asian countries (i.e., China, Korea) and were living in the Greater Toronto Area (GTA) for education and employment. Participants were right-handed, had normal or corrected-to-normal vision, and reported no history of neurocognitive disorders. Procedures were approved by the University of Toronto Research Ethics Board (protocol #44256). Participants provided informed consent and received monetary compensation.

We recruited East Asian participants because prior EEG work [15] found robust behavioral and decoding ORE in this population. Participants born and raised in East Asia also provided a contrast to individuals raised in a highly diverse multicultural region, such as the Greater Toronto Area, where attenuated ORE has been reported [7]. This design focuses on how viewing format changes race-related neural representations within the same observers.

Sample-size planning considered both behavioral ORE and its neural decoding counterpart for individual faces. We assumed a paired-samples effect size *d_z_* = .70 based on prior behavioral studies [43,62] and used .74 for the SR decoding advantage previously reported in East Asian participants [15]. A priori power analyses for two-tailed paired-samples t tests (α = .05, power = 80%; G*Power [63]) indicated minimum sample sizes of 19 and 17 participants, respectively. We therefore targeted at least 19 participants and retained 22.

### Stimuli

Stimulus generation relied on a StyleGAN2 architecture [64] trained from scratch to produce high-quality synthetic faces (see Supplementary Methods). Training relied on equally sized sets of East Asian and White faces, equally divided between male and female. This aimed to balance demographic representation in the training data, given reported race– and sex-related biases in generated faces [65,66].

Four subsets of 13 images, two per race, were selected to depict female-appearing faces with neutral expressions and frontal viewpoints (Fig. 1a), sampling different portions of the GAN latent space (Supplementary Methods; Supplementary Fig. 6). Low-level image properties were normalized; the sampling and pixel-distance assessment are detailed in Supplementary Methods.

Of the 13 faces in each sampling stratum, six were assigned to ensemble construction, in addition to being presented individually, while seven faces were presented only as individual-face stimuli (total N = 52 faces). Ensemble stimuli were constructed as: homogeneous (six same-race faces), outlier (five faces from one race and one from the other) and equal-split (ES; three faces per race) displays (Fig. 1b). Specifically, we constructed four homogeneous ensembles (i.e., one per sampling stratum across the two races), four outlier ensembles (by replacing one face from each homogeneous ensemble with one from the other race) and two ES ensembles (by combining three faces of each race from homogeneous ensembles). Ensemble faces were arranged in a roughly oval configuration around a central fixation cross. The spatial position of individual faces within each ensemble was randomly assigned on each experimental trial.

The GAN provides a continuous stimulus space, as used in related image-generation work [67], in which latent vectors can be averaged to define ensemble summary targets. Latent-space averaging generates an image from a mean vector, avoiding the feature misalignment of unregistered pixel averaging and other visual artifacts introduced by conventional morphing while providing high-fidelity estimates the group’s average appearance. We averaged constituent *W*-space vectors to obtain each model-defined ensemble mean and rendered it through the generator (Fig. 1b). Race-level average images were similarly generated from all *W* vectors used to define the sampling strata (Supplementary Methods).

In total, this procedure yielded 64 individual-face stimuli (13 faces per sampling stratum plus 10 ensemble summaries and 2 race averages). The remaining images from each stratum served as a validation set for the evaluation of reconstruction results.

### Experimental procedure

Participants first completed two versions of the CFMT, one with Chinese face stimuli [43] and one with White face stimuli [44]. These measures were used to (i) assess face-recognition proficiency relative to normative performance ranges [43,68] and (ii) quantify individual-level ORE behaviorally. Both CFMT versions have documented reliability and validity [43,44] and served to estimate ORE using a common subtraction approach (SR CFMT score minus OR CFMT score; [6,62]).

EEG testing comprised 32 blocks in one 2.5-hour session: 16 individual-face and 16 ensemble blocks presented in alternation. Participants were instructed to press the space bar when they detected male face oddballs, thus maintaining attention without an explicit identity, race or summary judgment (Fig. 1c). In each individual-face block, every stimulus appeared four times (totaling 256 trials), with an additional ∼10% of trials being male oddballs, totaling 280 trials. In each ensemble block, each of 10 distinct ensembles appeared 12 times (plus an additional 12 male oddball trials), to allow robust estimation of evoked responses across multiple repetitions. Presentation order was pseudorandomized to avoid consecutive target events and to reduce sequential dependencies.

Across blocks, stimuli were presented for 300 ms, followed by a variable 500-600 ms inter-stimulus interval during which a central fixation cross replaced the stimulus. Stimulus presentation and response collection were implemented in Psychtoolbox [69,70]. Stimuli were displayed on a 60 Hz monitor at 1920 × 1080 resolution, viewed from approximately 80 cm. Individual faces subtended approximately 3°×3° visual angle, whereas ensembles subtended approximately 10°×10° visual angle. The session began with two practice blocks (one for individual faces, one for ensembles) to familiarize participants with the male-oddball detection task.

### Statistics and Reproducibility

#### Statistical analyses

EEG and CFMT analyses relied on the same 22 participants while attribute validation used a separate sample of 44 participants. Tests were two-tailed with α = .05 unless a directional hypothesis was explicitly stated; correction families are specified below. Conventional parametric tests and standard Wilcoxon signed-rank tests were conducted in JASP 0.19.3. Attribute judgments were analyzed in R using ARTool 0.11.2 for aligned rank transform (ART) ANOVAs [71] and ART-C contrasts [72], and clusrank 1.0-4 [73] for clustered Wilcoxon signed-rank tests following the Rosner– Glynn–Lee procedure. Time-resolved decoding was evaluated using cluster-based permutation t-tests implemented in MNE-Python 1.11.0 [74]. Model specifications, permutation procedures, and multiple-comparison corrections are detailed in the corresponding analysis subsections.

Paired-samples *t* tests and repeated-measures ANOVAs evaluated mean differences and factorial effects, whereas Wilcoxon signed-rank tests evaluated chance performance, face-space distances and stand-alone reconstruction contrasts. For conventional repeated-measures ANOVAs, Shapiro–Wilk and, where applicable, Mauchly’s tests detected no significant departures from normality and sphericity, respectively. Comparisons of temporally cumulative decoding and reconstruction accuracy against chance used two-sided Wilcoxon signed-rank tests, applied consistently across conditions because some accuracy distributions departed from normality. The factorial ANOVAs directly tested whether the SR–OR difference varied with viewing format. ART ANOVAs evaluated factorial effects on bounded attribute-judgment proportions. Cohen’s *d* was standardized by the SD of paired differences for standalone paired *t* tests and by the pooled SD across the model’s condition cells for ANOVA post hoc contrasts; ANOVA effect sizes are partial *η*^2^.

#### Behavioral data analysis

Participants’ SR CFMT scores were inspected to verify that face-recognition ability fell within the expected normative range. The behavioral ORE was tested by comparing SR and OR CFMT scores using a two-tailed paired-samples *t*-test. Individual ORE magnitude was calculated as the SR CFMT score minus the OR CFMT score [6,62].

#### EEG acquisition and preprocessing

EEG was recorded using a 64-channel BioSemi ActiveTwo system (BioSemi B.V.), with electrodes positioned according to the international 10-20 system. Electrode offsets were maintained below 40 mV throughout acquisition. Signals were hardware low-pass filtered with a fifth-order sinc filter with a half-power cutoff of 204.8 Hz and digitized at 512 Hz with 24-bit resolution.

Offline preprocessing relied on Letswave 6 [75]. Continuous data were bandpass filtered using a zero-phase Butterworth filter (0.1-40 Hz, 24 dB/octave) and epoched from −100 to 900 ms. Epochs underwent DC removal, linear detrending, and baseline correction. Noisy channels were interpolated when necessary (maximum of two per participant) and the data were re-referenced to the common average. Ocular artifacts were attenuated using Infomax ICA [76,77].

Oddball sensitivity was near ceiling (mean *d*′ = 4.997). Male-target trials, false alarms, and trials with residual artifacts were excluded. We retained an average of 98.4% of trials per participant.

#### Temporally cumulative decoding

Decoding analyses focused on 12 occipitotemporal (OT) electrodes: P5, P7, P9, PO3, PO7, and O1 in the left hemisphere and P6, P8, P10, PO4, PO8, and O2 in the right hemisphere. These electrodes were selected because they reliably support temporal decoding of facial information [46, 78].

For all decoding approaches, pairwise classification was performed for both individual-face and ensemble stimuli using a linear support vector machine (C = 1) with leave-one-block-out cross-validation across 16 blocks. This produced a classification-accuracy estimate for every stimulus pair and participant. The overall decoding and reconstruction procedure is illustrated in Fig. 2 (face images in Figs. 2 and 6 are synthetic stimuli and EEG-based reconstructions generated using the StyleGAN2 model trained for this study; see Stimuli and Supplementary Methods).

Broad-window decoding, also termed temporally cumulative decoding in prior work [15], concatenated rather than averaged activity across time. Specifically, ERP amplitudes were concatenated across the 12 OT electrodes and all time points between 50 and 650 ms, yielding 3,684 features per observation (12 electrodes × 307 time points). This interval captures information distributed across early and later stages of face processing [35,46].

Within each block, repetitions of the same stimulus were averaged, and each electrode-by-time-point feature was z-scored. This yielded 16 observations per stimulus for each participant, separately for individual-face and ensemble blocks.

Individual-face accuracies were summarized for SR, OR, and between-race pairs. Classification accuracy was tested against 50% chance with two-tailed Wilcoxon signed-rank tests and Bonferroni correction across the three individual-face comparison types. Comparison types were evaluated with a one-way repeated-measures ANOVA and post hoc contrasts, Bonferroni-corrected across the three pairwise comparisons.

Ensemble decoding was summarized for homogeneous SR pairs, homogeneous OR pairs, between-race homogeneous pairs, ES ensemble comparisons, and outlier-ensemble comparisons. Separate one-way repeated-measures ANOVAs were conducted for homogeneous and ES ensembles, followed, where warranted, by post hoc contrasts with Bonferroni correction across the pairwise contrasts within the relevant omnibus analysis.

To test whether the SR-OR decoding difference varied across viewing formats, participant-level temporally cumulative accuracies were entered into a 2 × 2 repeated-measures ANOVA with viewing format (individual face, homogeneous ensemble) and stimulus race (SR, OR) as within-participant factors. The within-format comparisons described above characterized the direction of this interaction.

Constituent controls used individual-face EEG recordings restricted to the faces composing homogeneous ensembles. Pairwise accuracies were averaged separately for constituent pairs belonging to the same ensemble and pairs belonging to different ensembles of the same race. For each comparison, SR and OR accuracies were compared across participants using one-tailed paired t tests of the directional SR-over– OR hypothesis for within-ensemble and between-ensemble face pairs; the two reported *p* values were unadjusted.

#### Neural face space

To characterize the geometry revealed by temporally cumulative decoding, pairwise accuracies were arranged as representational dissimilarity matrices and submitted to classical metric MDS separately for individual faces and ensembles, with larger values/distances indicating higher discriminability. For each participant and stimulus type, we retained sufficient dimensions to account for ≥90% of the variance. Euclidean distances across all retained dimensions quantified representational separation.

For individual faces, pairwise distances in face space were computed across all retained dimensions and separately averaged for SR, OR, and between-race pairs. Then, distances were compared across different types of pairs using two-tailed Wilcoxon signed-rank tests across participants. Face spaces were also estimated from participant averages following the same procedure for visualization purposes.

#### Participant space

PCA was applied separately to participant decoding profiles for individual faces and ensembles. For individual faces, each participant was represented by a 2,016-element vector containing all pairwise accuracies among the 64 stimuli. Similarly, for ensembles, each participant was represented by a 45-element vector containing all pairwise accuracies among the 10 corresponding stimuli.

Scores on the first two principal components were correlated with SR CFMT performance and behavioral ORE magnitude using Pearson correlations, following prior evidence linking leading decoding components to these measures [15]. Components 3– 5 were examined exploratorily. Correlation *p* values were unadjusted.

Individual-face and ensemble participant configurations were compared via Procrustes alignment, with mean squared alignment error used to estimate the fit. Significance was evaluated by shuffling ensemble participant labels 10,000 times and recomputing the alignment and corresponding error. Mean SR and OR decoding accuracies were also correlated across formats to assess correspondence in overall performance.

#### Temporal dynamics

Pairwise decoding was repeated across ∼20 ms windows comprising 10 adjacent samples (∼1.95 ms each) and advanced one sample at a time from −100 to 900 ms to yield 503 overlapping windows [2,35].

Time-resolved decoding was evaluated using two-sided, one-sample cluster-based permutation t-tests [74] applied to participant-level decoding accuracies after subtracting permutation-derived chance estimates. Temporally adjacent samples exceeding a two-sided cluster-forming threshold of uncorrected *p* < .05 were grouped into separate positive and negative clusters. Cluster mass was defined as the sum of *t*-statistics within each cluster. Significance was assessed using 10,000 participant-wise sign-flip permutations and the maximum absolute cluster mass across time, controlling the cluster-level familywise error rate at .05.

The same procedure was applied to within-participant SR–OR differences between decoding time courses, separately for individual faces and homogeneous ensembles.

Additional analyses evaluated ES, SR-outlier and OR-outlier ensemble decoding.

To test whether race differences emerged at an intermediate temporal resolution, decoding also considered spatiotemporal patterns across five consecutive intervals spanning 100-650 ms: 100-210, 210-320, 320-430, 430-540, and 540-650 ms. Different decoding profiles were compared within each interval using Wilcoxon signed-rank tests with Bonferroni correction across the five windows.

#### Sensor-level searchlight decoding

Local sensor analyses used the same 50–650 ms interval; procedures and correction families are detailed in Supplementary Methods (Supplementary Figs. 4 and 5).

#### Reconstruction of individual-face and ensemble representations

To recover the representational content underlying EEG decoding, we conducted reconstruction in StyleGAN2’s *W* space. The procedure combined an established framework for image-based reconstruction from behavioral and neural data [42,45,46,79] with a GAN latent-vector reconstruction approach used in recent behavioral work [16] (Fig. 2; Supplementary Methods).

For each participant, we first estimated an individual-face neural space (see Neural face space). For each dimension, we derived a latent feature vector by weighting the face stimuli’s *W* vectors by their neural-space coordinates, analogous to reverse correlation. A target was projected into a neural space estimated without it, and its coordinates weighted a linear combination of the latent feature vectors. The generator rendered the estimated *W* vector as an image. For ensembles, cross-format decoding against individual faces supplied the relationships needed for projection into the individual-face neural embedding. Supplementary Methods detail the alignment and feature-estimation procedure.

The TO followed the same procedure using pairwise *W*-space distances instead of neural dissimilarities. Its reconstructions provided a model-derived reference for the selected stimulus set and for independent judgments of recovered attributes (Supplementary Methods).

Target-matching accuracy was the proportion of comparisons in which a reconstructed latent vector was closer to its target than to a held-out validation foil, using Euclidean distance in *W* space. Individual-face foils matched the target’s race, and ensemble foils matched the racial composition of the ensemble summary target (Supplementary Methods). Thus, 100% accuracy indicates closer matches to the target than to all validation foils, rather than complete recovery of latent or pixel information. Two-sided Wilcoxon signed-rank tests across participants assessed accuracy against 50% chance and between stimulus types.

Finally, to test whether the SR-OR reconstruction difference varied across viewing formats, participant-level reconstruction accuracies for SR and OR individual faces and homogeneous ensembles were additionally entered into a 2 × 2 repeated-measures ANOVA with viewing format (individual face, homogeneous ensemble) and stimulus race (SR, OR) as within-participant factors. ANOVA effect sizes were reported as partial 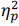; because race had two levels, its main effect is the SR-OR contrast averaged across viewing formats.

#### Behavioral validation of reconstructed attributes

A separate sample of 44 East Asian observers (age range: 18–35 years; 28 women) was recruited through Prolific (www.prolific.com). Procedures were approved by the University of Toronto Research Ethics Board (protocol #43487). Participants provided informed consent and received monetary compensation.

Two observers evaluated each of the 22 reconstruction sets, comprising images reconstructed from one EEG participant’s data and their TO counterparts. On each trial, an EEG reconstruction and its TO counterpart were presented side by side until observers selected which appeared younger, more positive, or more emotionally intense. Each attribute was assessed in a separate block, repeated with image positions reversed, yielding six blocks of 74 trials and two judgments per pair and attribute. The task included all ensemble types, but analyses focused on individual faces and homogeneous ensembles to match the race-by-format comparisons used for decoding and reconstruction.

Selection rates were calculated for each observer, attribute, and race-by-format condition, averaging across targets and repeated presentations. Responses were recoded so that values above 50% indicated that the EEG reconstruction appeared older, more positive, or more emotionally intense than its TO counterpart. These scores index relative appearance, rather than absolute attribute values or perceptual error.

Condition-wise selection rates were compared against 50% using two-tailed clustered Wilcoxon signed-rank tests, with reconstruction set as the clustering unit. Zero differences were excluded and significance was estimated using 100,000 random sign flips, applying a shared sign within each reconstruction set. Benjamini–Hochberg FDR correction covered the 12 tests (four conditions × three attributes).

Separate 2 × 2 mixed-effects ART ANOVAs tested race, viewing format, and their interaction for each attribute. Models retained all 44 observers nested within 22 reconstruction sets, with observer random intercepts and mutually uncorrelated set-level intercepts and slopes for race, format, and their interaction. Random slopes used centered race and format contrasts and their product. Type III Wald *F* tests used Kenward–Roger degrees of freedom. Two-sided ART-C contrasts used FDR correction across six comparisons per attribute. Reported *p*-values for condition-wise and ART-C tests are FDR-adjusted; omnibus *p*-values are unadjusted.

## Data availability

All data are publicly available through the Open Science Framework at https://osf.io/yvep6/.

## Code availability

Analysis scripts are available through the Open Science Framework at https://osf.io/yvep6/.

## Use of AI

ChatGPT (OpenAI) assisted with enhancing the readability of the manuscript, literature searches, and checks of statistical reporting. The authors critically reviewed and edited the output, verified the scientific content and references, and take full responsibility for the final manuscript.

## Supporting information

Supplemental Methods and Results

## Acknowledgements

This work was supported by a Discovery Grant from the Natural Sciences and Engineering Research Council of Canada (NSERC) to A.N.

## Author contributions

Conceptualization: M.S., M.A.S., J.S.C., A.N.; methodology: M.S., J.S.C., A.N.; software: M.S.; formal analysis: M.S., M.A.S.; investigation: M.S., H.N.J., Z.A., O.I.; data curation: M.S., H.N.J., Z.A., O.I.; visualization: M.S.; writing—original draft: M.S.; writing—review and editing: M.S., H.N.J., Z.A., O.I., J.S.C., A.N.; supervision: J.S.C., A.N.; project administration: J.S.C., A.N.; funding acquisition: A.N. All authors approved the final manuscript.

