## Supplemental Methods and Results for "Ensemble viewing reshapes neural representations of same-race and other-race faces"

#### *Supplementary Information*

Moaz Shoura, Han Ning Jiang, Zaynab Azeem, Otilia Iancu, Marco A. Sama, Jonathan S. Cant, and Adrian Nestor

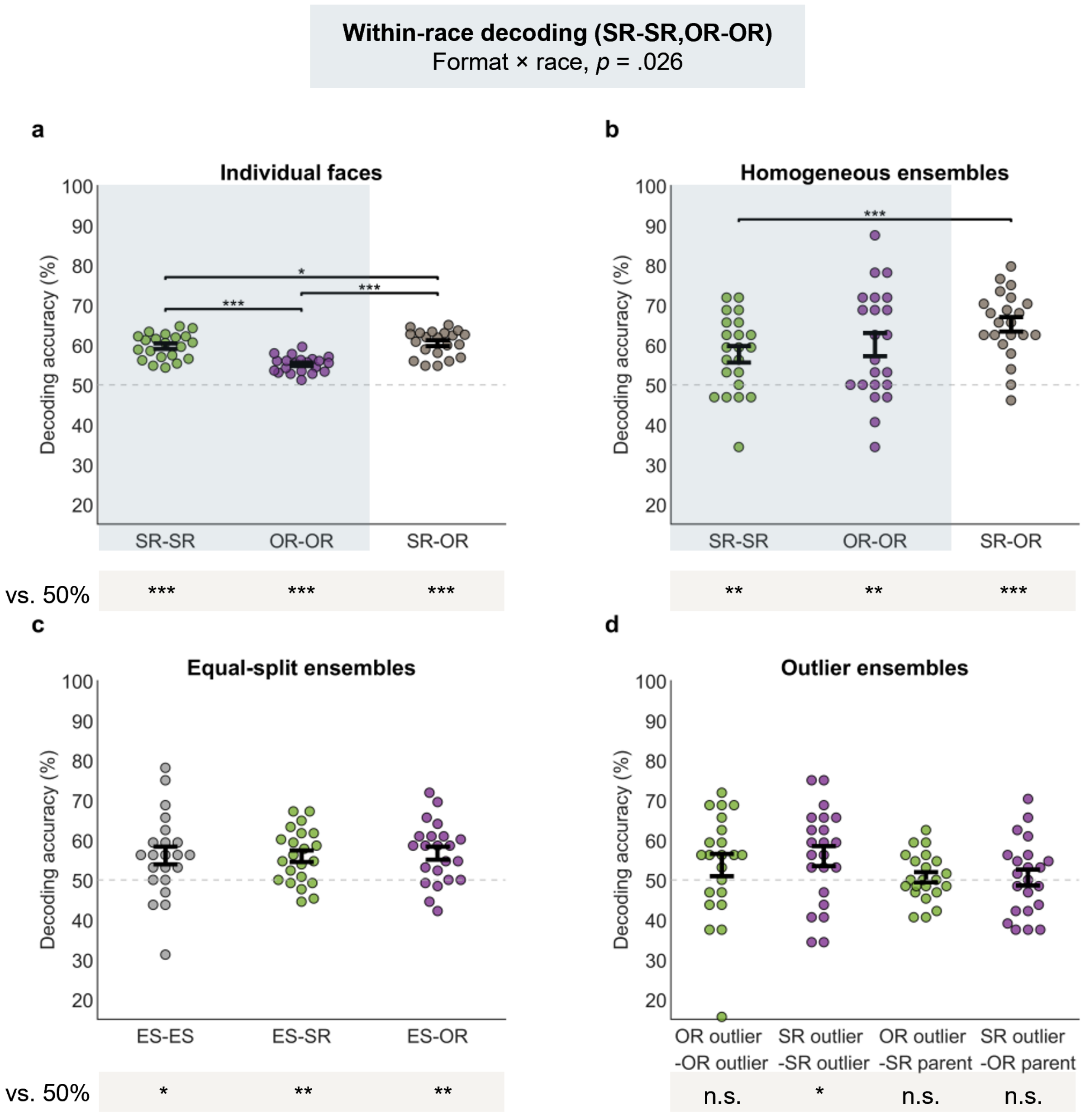


**Supplementary Fig. 1.** Broad-window decoding using all 64 electrodes over 50–650 ms. Accuracy values were generally lower than with the 12 occipitotemporal electrodes used in the main analysis, but the overall patterns were similar. **a** Individual-face decoding yielded a same-race (SR) advantage over other-race (OR) faces ($p < .001$), with between-race accuracy exceeding both SR ($p < .05$) and OR ($p < .001$) accuracy. **b** Between-race homogeneous-ensemble accuracy exceeded SR accuracy ($p < .001$). Pale-blue backgrounds in (**a**) and (**b**) identify the within-race categories entered into the viewing-format × race analysis; between-race decoding was excluded. **c** Equal-split (ES) ensembles were decoded above chance from one another and from homogeneous ensembles, without reliable differences among these three comparisons. **d** SR-outlier ensembles were decoded above chance from one another, but not from their parent homogeneous ensembles. OR-outlier ensembles were not decoded above chance from one another or from their parent homogeneous ensembles. Dots represent participants (*n*

= 22). Error bars indicate ± SEM. Dashed lines mark 50% chance. Brackets indicate two-sided post hoc contrasts with Bonferroni correction across three comparisons per panel in **(a)** and **(b)**. Grey strips show two-sided Wilcoxon signed-rank tests against chance, with Bonferroni correction across the three tests. † $p < .10$, * $p < .05$, ** $p < .01$, *** $p < .001$; *n*.s., not significant.

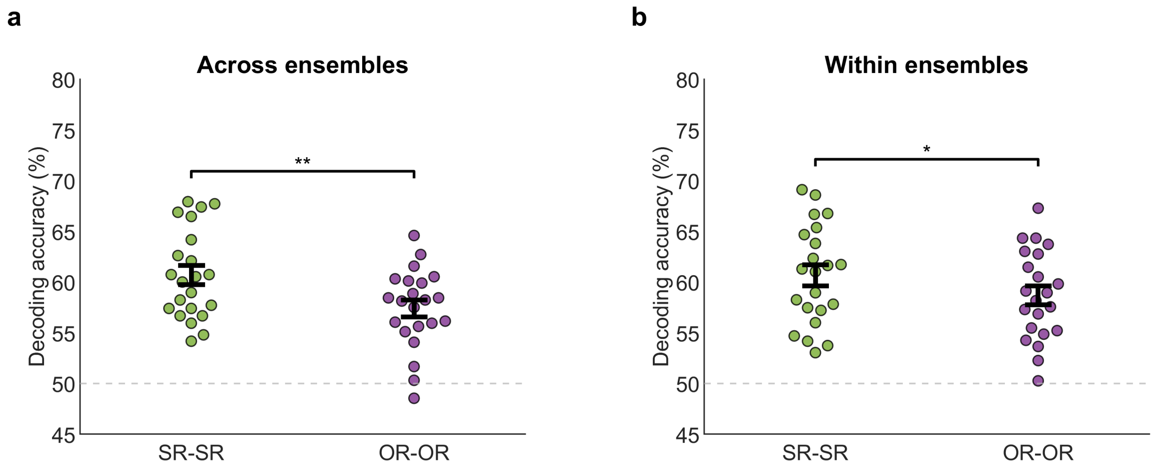


**Supplementary Fig. 2.** Broad-window decoding of individually presented ensemble constituents. Decoding used individual-face EEG responses restricted to the faces composing homogeneous ensembles. SR constituents were more discriminable than OR constituents both **a** for pairs drawn from different homogeneous ensembles of the same race and **b** for pairs within the same homogeneous ensemble (one-tailed paired-samples *t* tests across 22 participants; *p* values unadjusted across the two contrasts; * $p < .05$, ** $p < .01$).

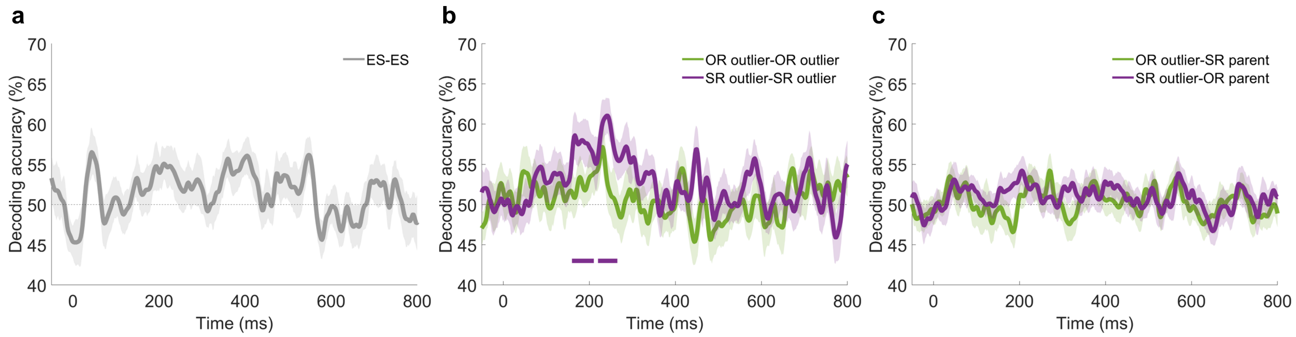


**Supplementary Fig. 3.** Time-resolved decoding of equal-split (ES) and outlier ensembles. **a** Decoding between the two ES ensembles. **b** Decoding within SR-outlier and OR-outlier types. Only SR-outlier decoding showed a cluster-corrected above-chance interval, at approximately 150–250 ms (purple bar). **c** Decoding outlier ensembles from their parent homogeneous ensembles, which shared five of six faces. No comparison in (**c**) yielded a corrected above-chance interval. $n = 22$ participants. Shading indicates ± SEM; horizontal bars denote cluster-corrected $p < .05$.

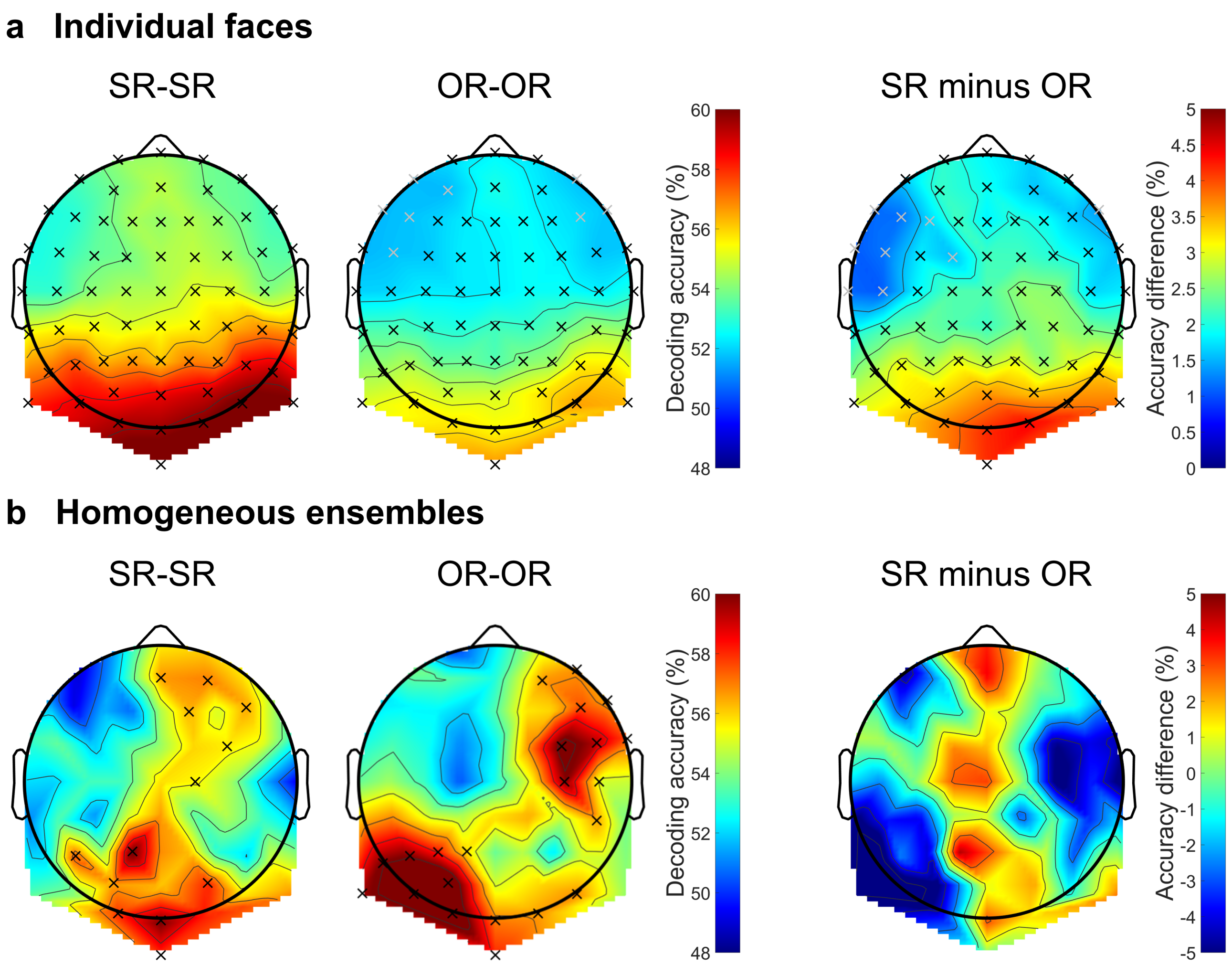


**Supplementary Fig. 4.** Sensor-level decoding reveals a widespread same-race advantage for individual faces. **a, b** Searchlight decoding of same-race (SR) and other-race (OR) individual faces (**a**) and homogeneous ensembles (**b**). Each searchlight combined activity from one electrode and its five nearest neighbors over 50–650 ms. The first two columns show mean within-race decoding accuracy; the final column shows SR minus OR accuracy. Crosses mark significant effects across 64 sensor neighborhoods (two-tailed Wilcoxon signed-rank tests, $p_{FDR} < .05$). Individual-face decoding exceeded 50% chance throughout the scalp and showed widespread SR advantages. Ensemble decoding exceeded chance in several neighborhoods, but no SR–OR difference survived correction. In the individual-face difference map, grey and black crosses indicate $p_{FDR} < .01$ and $p_{FDR} < .001$, respectively. $n$ = 22 participants.

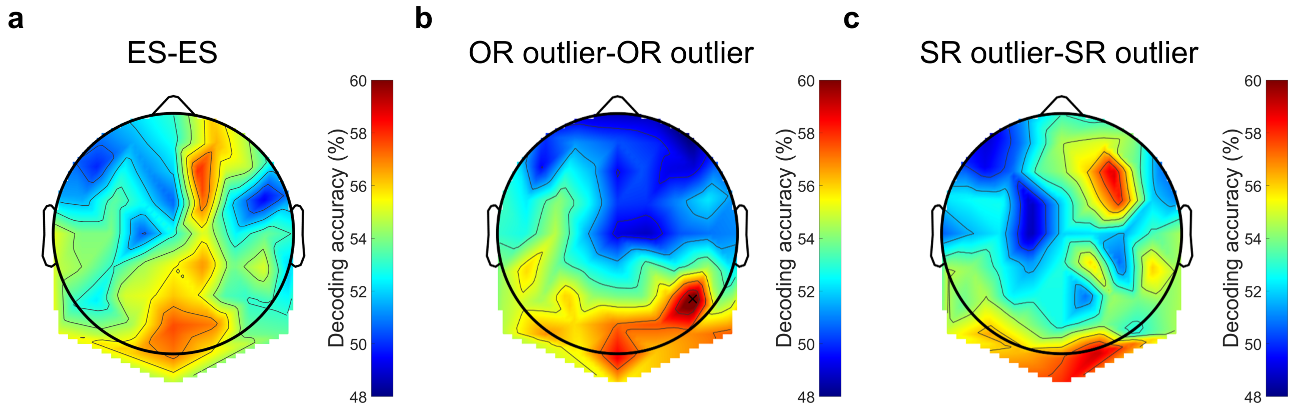


**Supplementary Fig. 5.** Sensor-level searchlight decoding for equal-split (ES) and outlier ensembles. Accuracy was estimated from each electrode and its five nearest neighbors over 50–650 ms. Panels show decoding between **a** ES, **b** OR-outlier, and **c** SR-outlier ensembles. The black cross in (**b**) marks the P6 neighborhood, the only searchlight with above-chance decoding after FDR correction across 64 neighborhoods ($p_{FDR} < .05$). Maps show mean accuracy across 22 participants.

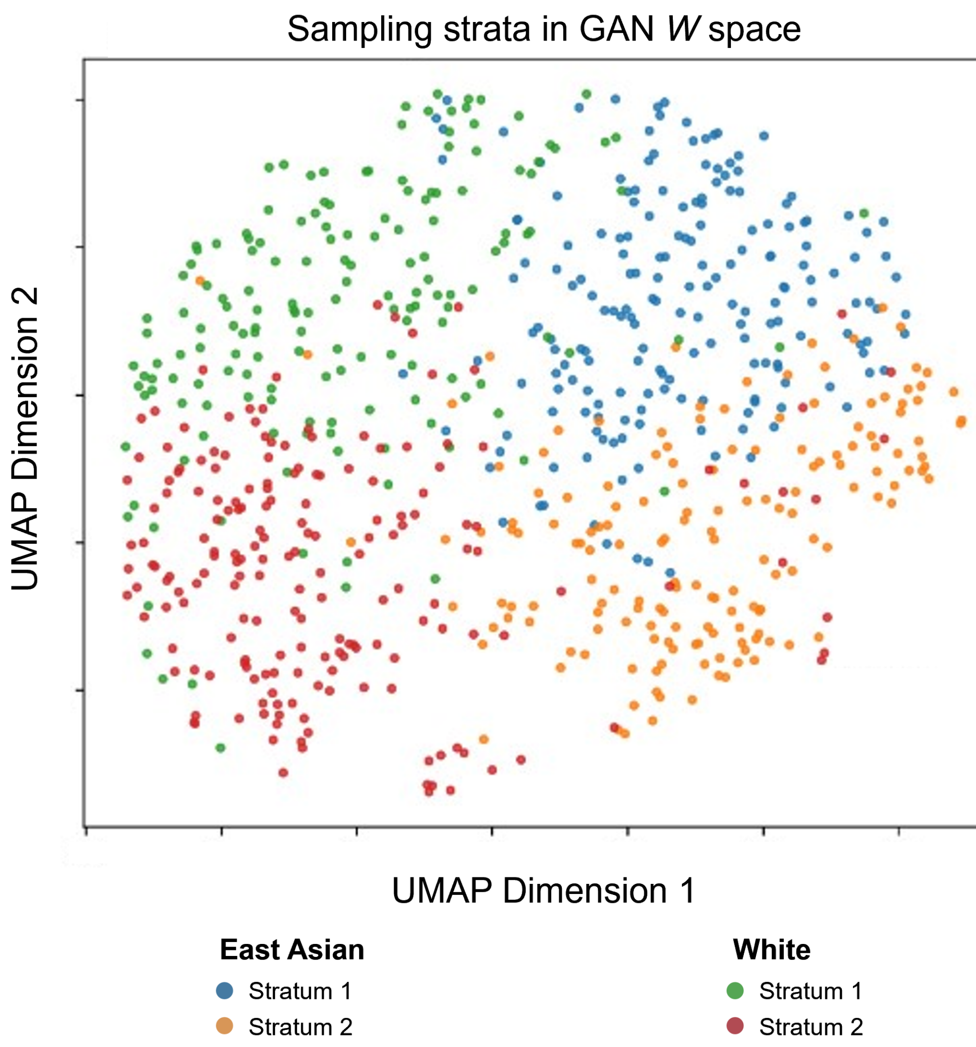


**Supplementary Fig. 6.** Descriptive UMAP visualization of race-specific sampling strata in StyleGAN2 *W* space. K-means ($k = 2$) was applied separately to East Asian-attributed and White-attributed face vectors in the original *W* space; colors indicate the two strata per race. UMAP provided the two-dimensional visualization and did not determine the sampling strata. Low silhouette scores for the two races (.031 and .037) indicate little separation between the strata.

### Supplementary Results

#### Supporting analyses of participant space

In the ensemble participant space, neither of the first two principal components was reliably associated with SR CFMT performance or behavioral ORE magnitude. Exploratory tests of components 3–5 were also nonsignificant.

Procrustes alignment identified correspondence between individual-face and ensemble participant configurations (main Results). In contrast, mean decoding accuracies were not significantly correlated across formats for either SR ($r$ = .119, $p$ = .597) or OR stimuli ($r$ = .357, $p$ =.103). Thus, the multivariate correspondence was detectable even though neither comparison of mean accuracy reached significance.

#### Sensor-level distribution of decoding information

Searchlight decoding characterized the scalp distribution of information combined over 50–650 ms using each electrode and its five nearest neighbors**.** Because scalp voltage is spatially mixed and adjacent searchlights overlap, the maps identify sensor-level information rather than localized cortical generators.

SR and OR individual faces were decoded above chance across all sensor neighborhoods (Wilcoxon tests, $p_{FDR} < .05$; Supplementary Fig. 4a), with widespread SR-over-OR differences. This agrees with the occipitotemporal (OT) and all-electrode broad-window analyses (Supplementary Fig. 1).

For homogeneous ensembles, above-chance SR and OR decoding involved overlapping posterior neighborhoods, including left occipitotemporal sites, as well as right-anterior neighborhoods (Supplementary Fig. 4b). No SR–OR difference survived FDR correction at any neighborhood (all $p_{FDR}$ values > .05). ES and outlier decoding also did not survive correction at any neighborhood except P6 for OR-outlier decoding ($p_{FDR} < .05$; Supplementary Fig. 5).

#### Supplementary Methods

##### StyleGAN2 training and image selection

We trained a StyleGAN2 generator [1] from scratch using 60,000 real face images, with horizontal-flip augmentation. The training set contained four equally sized race-by-sex subsets (15,000 images each). White-attributed faces were sampled from CelebA [2]; East Asian-attributed faces were sampled from CelebA, FairFace [3] and AFAD [4]. Images meeting a minimum resolution of 200 × 200 pixels were retained and resized to 256 × 256 pixels before training.

Training used StyleGAN2 configuration F, non-saturating logistic adversarial loss with R1 regularization and path-length regularization. Training continued until 10,000 kimg. Checkpoints were selected using Fréchet Inception Distance (FID; [5]) against a held-out subset of the curated corpus and visual inspection. The selected model achieved an FID of 3.21 for this reference distribution. Visual inspection complemented FID because the metric depends on the evaluation setup and need not track perceived image quality [6].

After training, we sampled 10,000 synthetic faces (256 × 256 pixels) at truncation $\psi = 0.5$ to balance image fidelity against identity diversity. DeepFace facial-attribute screening [7] retained faces classified as female and neutral in expression from both racial categories. Two human validators confirmed these attributes and excluded images with visible artifacts, nonfrontal views, or occlusions such as glasses, yielding 352 images per race. Demographic labels refer to perceived or attributed appearance rather than biological categories.

To sample different portions of each race-specific latent distribution, we applied $k$-means ($k = 2$) separately to the 352 $W$-space vectors per race. We ran 1,000 random initializations and retained the solution with the highest mean silhouette score for each race. The low scores (0.031 for East Asian and 0.037 for White face vectors) provided little evidence of well-separated groups. We therefore refer to these operational partitions as sampling strata (Supplementary Fig. 6).

From each of the four sampling strata (two per race), we selected 13 faces pseudorandomly, subject to no significant race difference in theoretical-observer target-matching accuracy (main Methods, Reconstruction of individual-face and ensemble representations; Fig. 6a). Images were normalized to common channel-wise means and contrasts in CIE L*a*b* space and projected back into $W$ space using StyleGAN2's standard gradient-based inversion procedure (1,000 steps; Adam optimizer $\eta_{max} = 0.1$). Projection quality was assessed with Learned Perceptual Image Patch Similarity (LPIPS) loss [8], with values of 0.029 and 0.032 for East Asian-attributed and White-attributed faces, respectively. Pairwise pixel-wise L2 distances were assessed across the four strata (one-way ANOVA, $p = .474$). This nonsignificant comparison does not establish equivalence of image properties.

**Image reconstruction procedure and theoretical observer**

A theoretical observer (TO) supplied reference reconstructions from the GAN's latent-space geometry using a procedure adapted from previous work [9,10,11]. We computed Euclidean distances between all pairs of individual-face $W$ vectors. For each individual-face target $k$, its row and column were omitted from the dissimilarity matrix used to derive reconstruction features. Metric multidimensional scaling (MDS) of this target-excluded matrix yielded a TO embedding, retaining up to 40 dimensions that accounted for ≥90% of the variance.

Coordinates along each dimension were z-scored. A latent feature vector was derived for each dimension by averaging non-target $W$ vectors weighted by these coordinates, analogous to reverse correlation. A space A′ containing the target and reference faces was aligned to the target-excluded space A by Procrustes transformation, which projected the target into A. Target-to-reference dissimilarities thus supplied the target's coordinates, while its $W$ vector was excluded from feature derivation. Feature vectors were combined linearly, weighted by the target's coordinates, to estimate its $W$ vector, which the generator rendered as an image.

EEG reconstruction followed the same framework, replacing GAN latent-space distances with participant-specific pairwise decoding accuracies to derive a neural face-space embedding. Ensemble representations were projected into this individual-face neural space using ensemble-to-individual-face decoding accuracies, then reconstructed from the corresponding latent feature vectors.

Target-matching accuracy was the proportion of comparisons in which a reconstructed $W$ vector was closer to its target than to a validation foil. Individual-face reconstructions were evaluated against same-race foils using Euclidean distance in $W$ space. Because model-defined ensemble means average multiple latent vectors, their variability can differ from that of individual faces. We therefore constructed composition-matched validation sets for evaluating ensemble summary targets. For SR homogeneous ensembles, we sampled sets of six vectors with replacement from the SR validation pool and averaged each set to generate 1,000 ensemble foil means. Foils for other ensemble types were generated analogously, preserving their racial composition (for example, three vectors per race for ES ensembles).

**Sensor-level searchlight decoding**

To characterize the scalp distribution of discriminative information, an electrode-wise searchlight analysis was conducted separately for individual faces and ensembles. Classification was performed at each electrode using that electrode along with its five nearest neighbors. The searchlight analysis used the same temporally cumulative 50-650 ms interval as the main decoding analysis.

Accuracy was tested against 50% chance at each electrode neighborhood using Wilcoxon signed-rank tests with false-discovery-rate (FDR) correction across 64 searchlights ($p_{FDR} < .05$). SR and OR individual-face and homogeneous ensemble decoding were also directly compared at each neighborhood.
